# Proteogenomics Reveals how Metastatic Melanoma Modulates the Immune System to Allow Immune Evasion

**DOI:** 10.1101/2021.04.10.439245

**Authors:** Jeovanis Gil, Yonghyo Kim, Beáta Szeitz, Viktória Doma, Uğur Çakır, Natália Pinto de Almeida, Yanick Paco Hagemeijer, Victor Guryev, Jenny G Johansson, Yogita Sharma, Indira Pla Parada, Zsolt Horvath, Jéssica de Siqueira Guedes, Gustavo Monnerat, Gabriel Reis Alves Carneiro, Fábio CS Nogueira, Boram Lee, Henriett Oskolas, Enikő Kuroli, Judit Hársing, Yutaka Sugihara, Magdalena Kuras, Roger Appelqvist, Elisabet Wieslander, Gilberto B Domont, Bo Baldetorp, Runyu Hong, Gergely Huszty, Laura Vizkeleti, József Tímár, David Fenyö, Lazaro Hiram Betancourt, Johan Jakobsson, Johan Malm, Aniel Sanchez, A. Marcell Szász, Peter Horvatovich, Melinda Rezeli, Sarolta Kárpáti, György Marko-Varga

## Abstract

Malignant melanoma (MM) develops from the melanocytes and in its advanced stage is the most aggressive type of skin cancer. Here we report a comprehensive analysis on a prospective cohort study, including non-tumor, primary and metastasis tissues (n=77) with the corresponding plasma samples (n=56) from patients with malignant melanoma. The tumors and surrounding tissues were characterized with a combination of high-throughput analyses including quantitative proteomics, phosphoproteomics, acetylomics, and whole exome sequencing (WES) combined with in-depth histopathology analysis. Melanoma cell proliferation highly correlates with dysregulation at the proteome, at the posttranslational- and at the transcriptome level. Some of the changes were also verified in the plasma proteome. The metabolic reprogramming in melanoma includes upregulation of the glycolysis and the oxidative phosphorylation, and an increase in glutamine consumption, while downregulated proteins involved in the degradation of amino acids, fatty acids, and the extracellular matrix (ECM) receptor interaction. The pathways most dysregulated in MM including the MAP kinases-, the PI3K-AKT signaling, and the calcium homeostasis, are among the most affected by mutations, thus, dysregulation in these pathways can be manifested as drivers in melanoma development and progression.

The phosphoproteome analysis combined with target-based prediction mapped 75% of the human kinome. Melanoma cell proliferation was driven by two key factors: i) metabolic reprogramming leading to upregulation of the glycolysis and oxidative phosphorylation, supported by HIF-1 signaling pathway and mitochondrial translation; and ii) a dysregulation of the immune system response, which was mirrored by immune system processes in the plasma proteome. Regulation of the melanoma acetylome and expression of deacetylase enzymes discriminated between groups based on tissue origin and proliferation, indicating a way to guide the successful use of HDAC inhibitors in melanoma. The disease progression toward metastasis is driven by the downregulation of the immune system response, including MHC class I and II, which allows tumors to evade immune surveillance. Altogether, new evidence is provided at different molecular levels to allow improved understanding of the melanoma progression, ultimately contributing to better treatment strategies.

**TOC figure:** 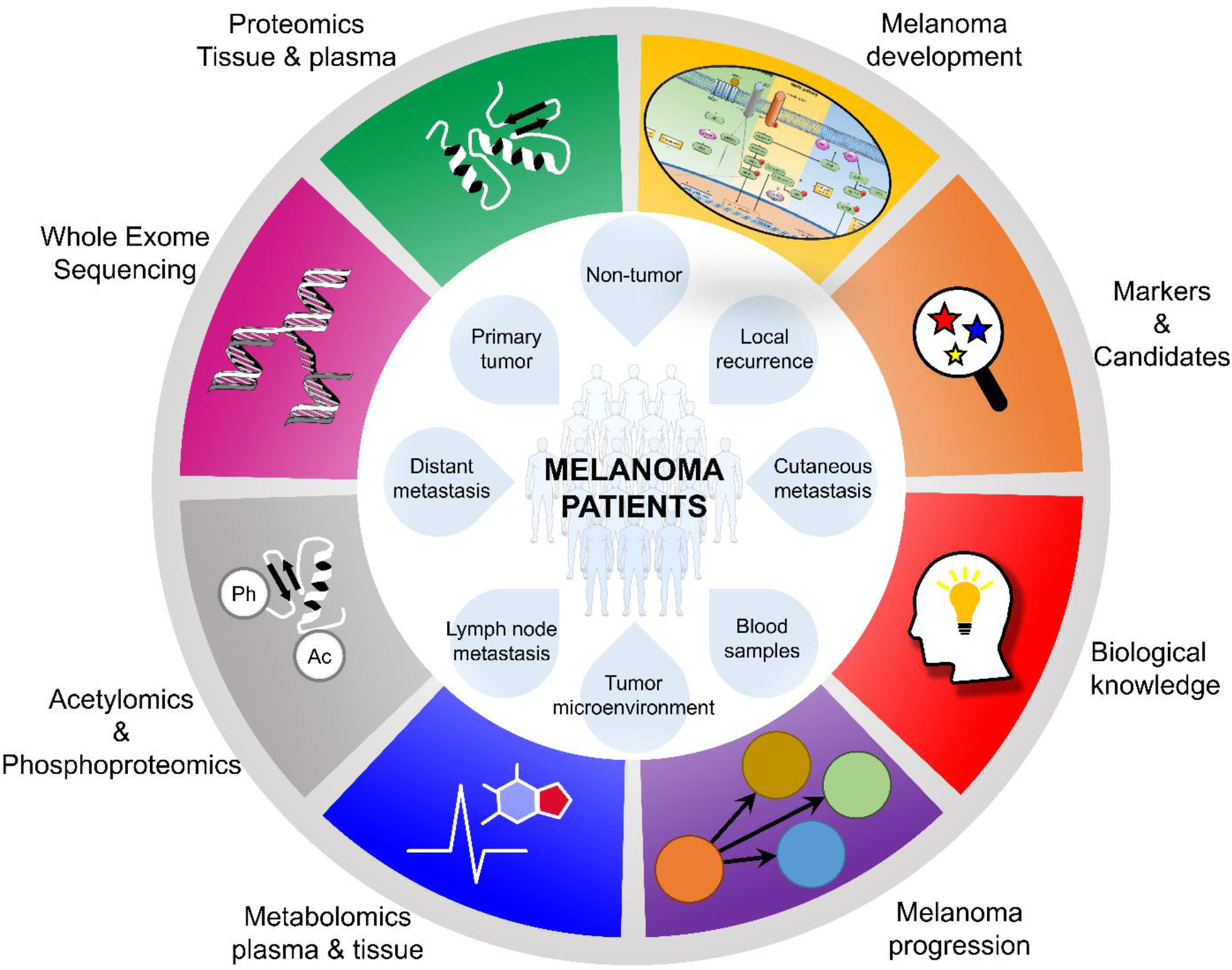

## 1. Introduction

Among the skin cancers, malignant melanoma is the deadliest, and its incidence and mortality are rising, particularly in Europe ^1,2^. However, the early surgical intervention, and more recently the advent of targeted therapy and immunotherapy have substantially improved the mortality rate of the disease. Melanoma is one of the most heterogeneous cancers at the genomic level, which results in a variety of distinct phenotypes of the disease ^3^. The high level of heterogeneity in melanoma has been previously linked to disease progression and short survival ^4,5^. Additionally, the levels of mutated proteins and dysregulation of their biological pathways, critically impact the tumor biology and survival ^6^. Signaling pathways such as the PI3K-AKT, mTOR, and MAPK are usually dysregulated in melanoma. BRAF, which is a kinase involved in the MAPK/ERK signaling pathway is the most frequently observed driver mutation in melanoma counting for approximately 50% of cases ^7^. Thus, a common treatment strategy to target this pathway is to use a combination of BRAF and MEK inhibitors. After a progression free period, however, most of the responsive patients in stage 4 of the disease develop resistance to treatment and ultimately die.

Large cohorts of melanoma samples have been studied through genomics, transcriptomics, proteomics in attempts to decipher the molecular mechanisms driving the development and progression of the disease ^8,9^. Phosphoproteomics and acetylomics on melanoma have shown dysregulation during the progression of the disease at the posttranslational level ^10,11^. Additionally, a recent international milestone was reported; the high-stringency blueprint of the human proteome ^12^, mapping the expression and functional annotation of the coded genome. In an attempt to integrate several high-throughput analyses into the biological understanding of the development and progression of melanoma, we performed a comprehensive multi-omic analysis including proteomics, PTM profiling, metabolomics, and whole-exome sequencing of 77 tissue samples coming from 47 melanoma patients, supplemented by the proteomic analysis of 56 plasma samples. The metabolic changes that take place during the development and progression of the disease were traced at different molecular levels. We demonstrated that the characterization of melanoma in such an integrative manner at different molecular levels led to a better biological understanding of the disease that will ultimately help to advance future therapeutic interventions

## 2. Results

### 2.1. Characteristics of the melanoma-related prospective cohort study

A cohort of 77 solid tissues, and 56 blood samples, originating from 47 different melanoma patients, enrolled in a prospective study, were submitted to a multi-omic analysis. Fresh frozen biopsies originated from different organs and body localizations, including non-tumors (**NT**; >2 cm from the tumor edges; n=11), tumor microenvironments (**TM**; tumor edge without countable tumor cells; n=6), primary tumors (**PT**; n=16), local recurrences (**LR**; n=6), cutaneous metastases (CM; n=10), regional lymph node metastases (**LN**; n=23), and distant metastases (**DM**; coming from gallbladder, brain, liver, spleen, and breast; n=5) (**Figure 1A**). Blood sampling was performed prior to the surgical isolation of the tumor tissues. The relevant clinical information of the patients included in the study is presented in **Table S1**. All samples were submitted to quantitative proteomics. Besides, the tissue samples were subjected to quantitative phosphoproteomics and lysine acetylation stoichiometry analysis (**Figure 1B**). Complementary whole-exome sequencing (WES) was performed on the tumor tissue samples (60). The disease presentation within the tumors from the patients is outlined in **Figure 1C**, where grading was made according to the immune competence, as well as the tumor burden encompassing the metastasis types, the primary and non-tumors.

**Figure 1.**
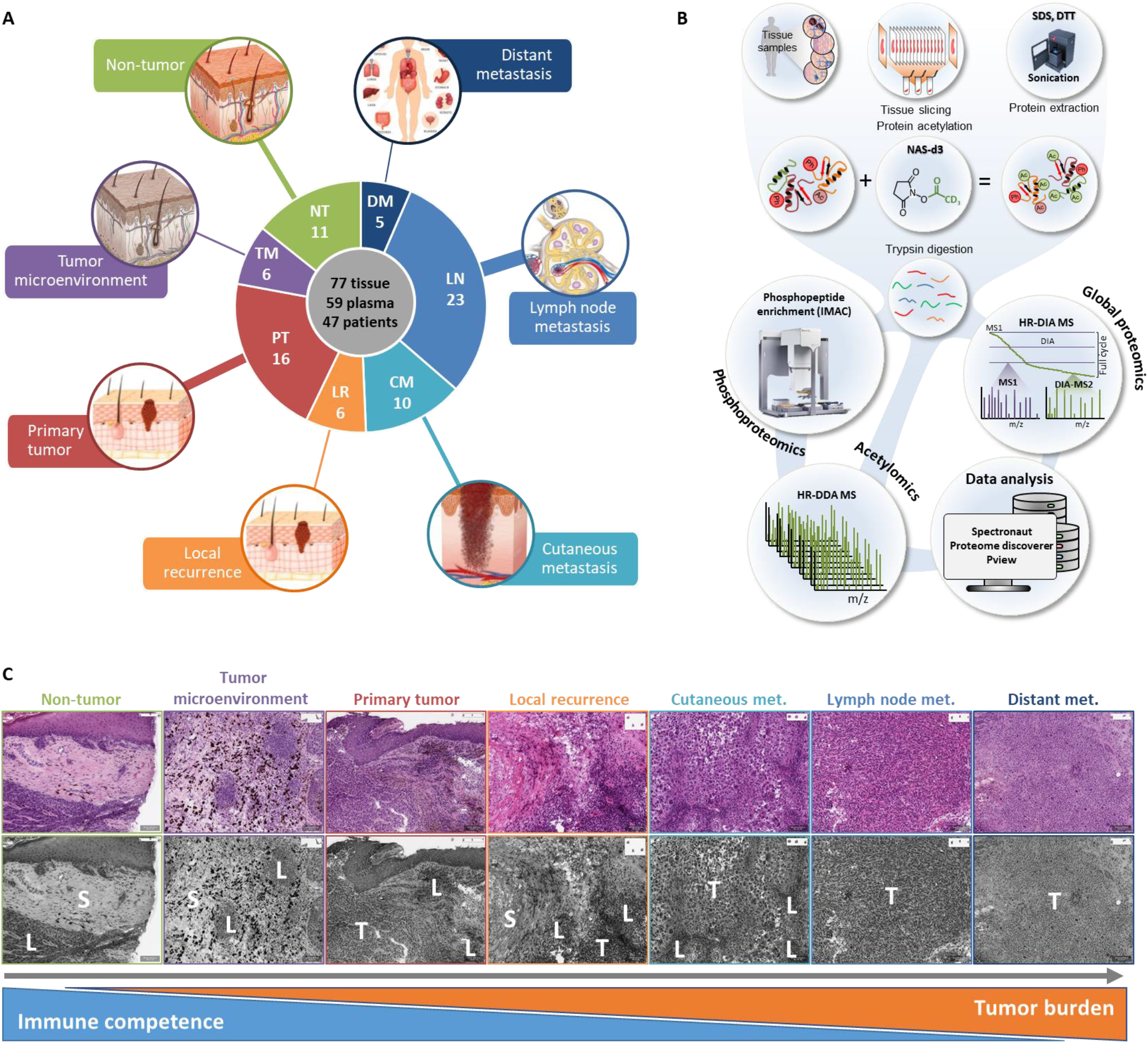
Melanoma-related cohort submitted to multi-omics approach. **A)** Tissue samples originated from non-tumor (green, n=11), tumor microenvironment (purple, n=6), primary tumor (n=16), local recurrence (orange, n=6), cutaneous metastasis (n=10), lymph node metastases (n=23), and distant metastasis (n=5). **B)** Workflow for the global proteomics, phosphoproteomics, and acetylomics used in tissue-derived samples. Tissue samples were sectioned (10 µM thickness), sonicated using Bioruptor plus in SDS containing buffer. The proteins were acetylated with N-acetoxysuccinimide-d3 (NAS-d3), followed by trypsin digestion. Generated peptides were divided into three different analyses; one fraction was directly injected onto LC-MS/MS system and analyzed following a data-independent analysis (DIA) method. A second fraction was submitted to an automated phosphopeptide enrichment analysis before LC-MS/MS analysis in a classical DDA method. The third fraction was measured following a DDA method without additional processing. Global proteomics data were analyzed with Spectronaut software, phosphoproteomics data was processed with Proteome Discoverer and acetylomics data by the combination of Proteome Discoverer and Pview. **C)** Spectrum of tumor-immune system imbalance is depicted during the progression of the disease. Starting from a relatively competent immunologic state both in peritumoral and intratumoral regions of low tumor load the pathological process ultimately results in high tumor burden with an exhausted immune system. Greyscale images correspond to the color figures above displaying areas of tumor (T), lymphoid infiltrate (L), and stroma (S). Oftentimes, this latter microenvironment adjacent to actual tumor cell nests is also dark (macroscopically), contains macrophages performing melanin phagocytosis.

### 2.2. Dysregulated pathways in melanoma carry a large mutation load

The tumor tissue samples were submitted to whole-exome sequencing (WES) analysis. In the absence of normal tissues as control, we relied on the gnomAD database’s non The Cancer Genome Atlas (TCGA) subset of ExAC release 1 to estimate the “population background”. The analysis revealed that melanoma does carry a higher number of somatic mutations, as compared to germline mutations (A>T and T>A SNP mutations, and indels are 15 and 224 times more abundant). However, the variants corresponding to SNP changes; A>G and T>C, were found to be underrepresented, whereas C>T and G>A were at comparable levels in somatic compared to germline mutations (**Table S2**). The relative ratio of different equivalent base-pair changes in somatic and germline SNPs variants are consistent across samples and the somatic patterns are different from the germline one. Germline patterns show high similarity with the population background except for A>G and T>C mutations, which is considerably larger in the germline pattern of the analyzed tumors compared to the population background (**Figure S1A-B**). The ratio of indels and SNPs in somatic mutations is substantially lower than in germline variants (0.81±0.03 and 74.53±4.48 SNPs/indels ratio respectively), indicating that melanoma is driven by indels rather than SNPs **(Figure 2D)**. Differences in this ratio were not observable, based on the sample origin. We also found that indels are slightly extending exon and protein sequences in somatic variants, while germline indels tend to mainly shorten them (**Figure S2**). The highly reproducible somatic patterns suggest that mutations in melanoma are driven by similar underlying molecular mechanisms across all patients.

**Figure 2.**
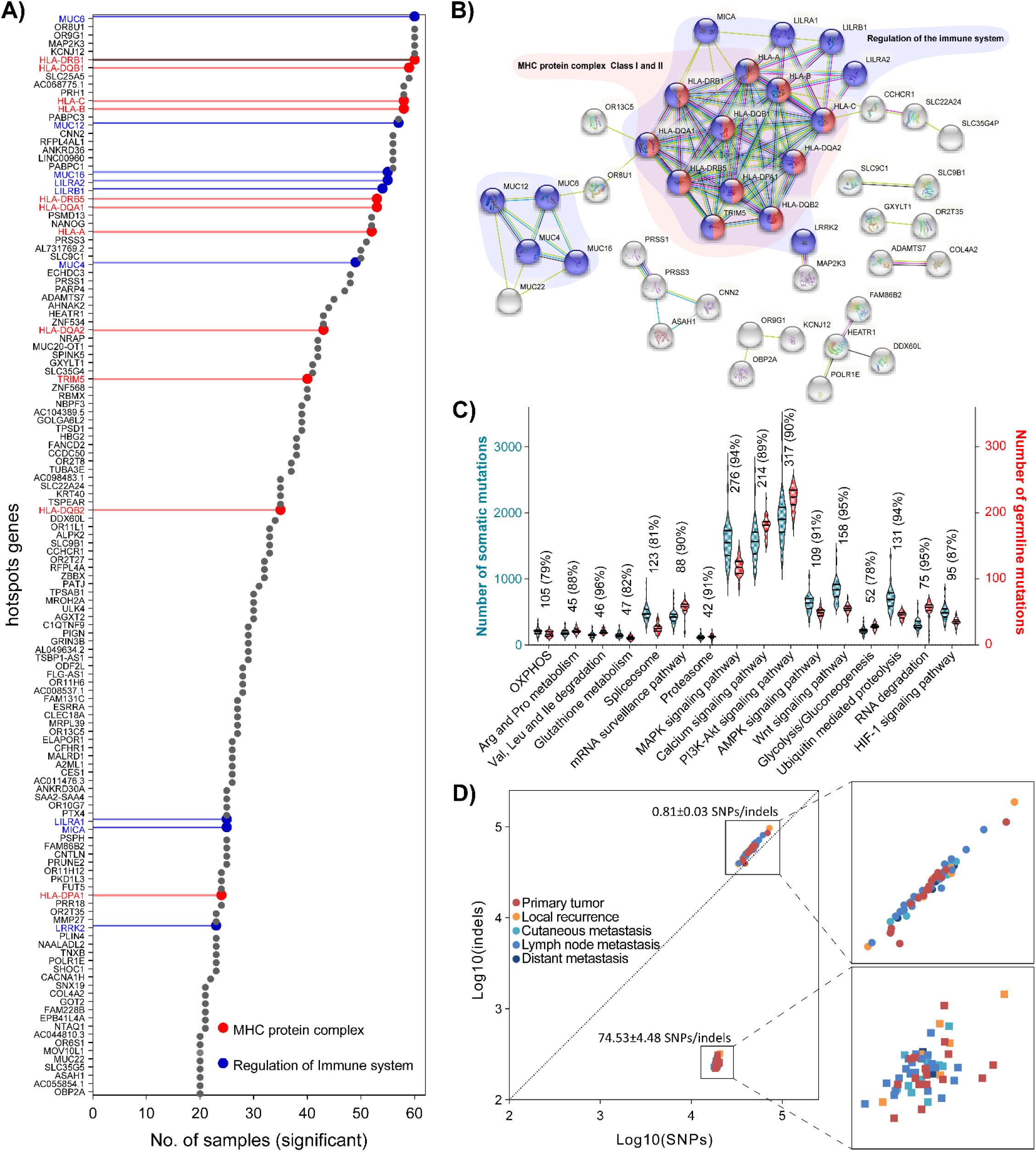
Most mutated genes in melanoma are involved in the regulation of the immune system. **A)** List of genes identified as hot spots in at least one-third (20 samples out of 60) of the tumor cohort. **B)** Protein-protein interaction network of the hot spot genes. Highlighted in red are the protein elements of the complexes MHC class I and II, and in blue additional proteins involved in processes related to the regulation of the immune system. **C)** Violin plots showing the number of somatic (left axis) and germline (right axis) mutations in genes involved in 16 KEGG pathways known to play crucial roles and found dysregulated in melanoma. The numbers inside the graph represent the number of mutations in the pathway and the percentage indicates the total number of genes detected in WES data relative to the total number of genes in the pathway. Data with the summary of the figure are outlined in **Table S4. D)** Number of mutations (indels vs. SNPs) for somatic and germline mutations in each sample on a logarithmic scale. Zoomed-in regions for somatic and germline mutations are shown.

The high genetic heterogeneity of melanoma tumors was highlighted by lower sequence coverage of somatic compared to germline variants. While germline mutations are expected to be present in all cells, somatic mutations are only present in tumor cells, and different clones carry different somatic mutations (**Figure S3**). We performed hot spot analysis to identify the genes most frequently found carrying somatic mutations across the samples using a Chi-squared test followed by a multiple testing with Bonferroni correction. We were able to locate the chromosomal regions with somatic mutation burden (**Figure S4**). Genes that were significantly mutated in at least one-third of the samples (20 out of 60) were selected as hot spots (132 genes in total) (**Figure 2A, Table S3**). The functional annotation analysis revealed that to a large extent, these genes are involved in the interaction and regulation of the immune system (**Figure 2B**). Particularly, the genes coding for the protein elements of both MHC complexes class I and II carry mutations in most melanoma samples. To a lesser extent, genes functionally related to metabolism and signaling cascades were also among the most mutated genes.

The most dysregulated pathways in melanoma found in this study at the transcriptional, translational, and posttranslational level, were also found to carry a significantly higher level of both germline and somatic mutations than the background population (**Figure 2C, Table S4**). In this sense, the pathways most affected by mutations in melanoma are the signaling cascades including MAP kinases and PI3K-AKT, and the calcium homeostasis. Not surprisingly, dysregulations of these pathways have been considered drivers in melanoma development and progression. On average, these pathways carry at least 11 times more somatic than germline mutations, which indicates that the genes of these pathways are heavily affected by the mutation burden in melanoma.

### 2.3. Proteome profiling of biopsies from melanoma patients

Tissue samples were submitted to a workflow that allows, from the same sample processing, to perform quantitative proteomic, phosphoproteomic, and lysine acetylation stoichiometry analysis ^10,13^ (**Figure 1B**). Briefly, the frozen tissue was sliced, and 15 consecutive slices were used for proteomics and PTMs analysis, or whole-exome sequencing. The first and last slices were H&E stained for computing the sample composition (tumor, stroma, and immune cell content). Proteins extracted, followed by chemically acetylated with NAS-d3 and digested with trypsin. Generated peptides were delimited by arginine residues. For global proteomics, peptides were directly analyzed by high-resolution mass spectrometry following a data-independent analysis (DIA) method. The phosphoproteome analysis included an automated phosphopeptide enrichment step using IMAC (Fe(III)-NTA), followed by high-resolution mass spectrometry data-dependent analysis (DDA). For lysine acetylation stoichiometry analysis, peptides were analyzed following a DDA method for acquisition (**Figure 1B**).

The proteomic analysis resulted in the identification and quantification of 9040 proteins with more than 2500 identified in all analyzed samples (**Figure 3**). On average, we identified approximately 7000 proteins in the tumor samples. 17237 phosphorylated peptides were identified in phosphoproteomics data, approximately 8000 in each sample. Additionally, we identified and determined the site-specific occupancy of 6844 acetylated peptides in at least 3 different samples, which correspond to 3024 different proteins. The plasma proteome analysis of the patients resulted in the identification of 1370 proteins.

**Figure 3.**
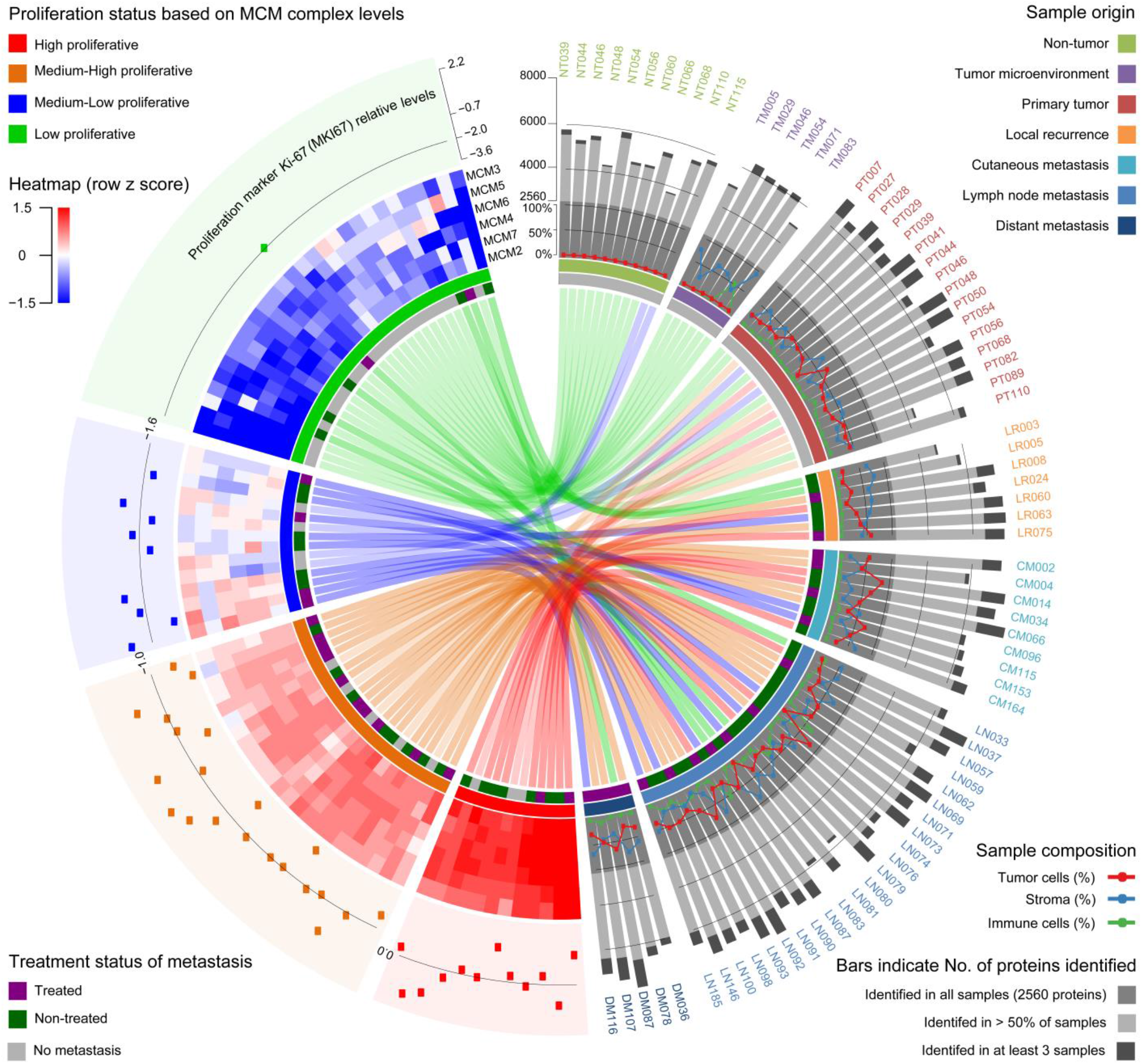
Proteomics of fresh frozen tissue biopsies from melanoma patients. A total of 77 samples coming from 47 different patients are grouped according to their origin. Eleven Non-tumor (NT) biopsies from more than 2 cm from the tumor boundaries, six tumor microenvironment (TM) samples taken from the tumor edges without the presence of tumor cells, sixteen biopsies from primary tumors (PT), six biopsies from local recurrence (LR) of the disease, ten biopsies of cutaneous metastases (CM), twenty-three metastases in regional lymph nodes, and five distant metastases (DM) coming from the gallbladder (DM036), brain (DM078), liver (DM087), spleen (DM107) and breast (DM116). Bars indicate the number of proteins quantified in each sample. The cell composition of each sample, taking into account tumor cell, stroma, and immune cells was estimated and expressed in percentage of the total sample area. The heatmap was built with the relative abundance levels shown as Z-scores of the six protein elements of the mini-chromosome maintenance complex and four clusters were created to represent the proliferation status of the samples. The abundance of the proliferation marker Ki-67 was plotted for each sample in the proliferation groups. A line indicates the mean value for each group relative to the most proliferative sample group. The colored connection arcs link the samples’ origin with their proliferation status.

From the proteomic data, the relative abundance profiles of the protein elements of the minichromosome maintenance (MCM) complex were extracted and submitted to hierarchical clustering analysis. As a resulting outcome, all samples were grouped into four groups from lower to higher levels of the complex elements (**Figure 3**). MCM is a hetero-hexameric complex required for DNA replication, which possesses DNA helicase activity ^14–17^. Due to its role in DNA replication and because their protein elements have been reported as proliferation markers and associated with progression in many cancers ^14,15,25,26,16,18–24^, we procured their protein abundance profiles as a measure of the proliferation-rate of the sample. The levels of the well-known proliferation marker Ki-67 showed significant differences between the proliferation groups, where the groups with higher MCM complex levels, also upregulate Ki-67 (**Figure 3**). As expected, non-tumor and tumor microenvironment samples were grouped in the low proliferative groups and Ki-67 was not detectable in any of them.

### 2.4. Melanoma proliferation relies on repressing the immune system response-related pathways

We sought to gain a deeper understanding of what processes are altered with increased tumor cell proliferation. This was addressed using the proteomic data of the 60 tumor tissue samples and 56 paired plasma samples as well as the Skin Cutaneous Melanoma TCGA PanCancer data with 443 tumor tissue samples (involving 20531 transcripts). TCGA tissue samples were classified into proliferation groups based on the MCM complex levels (in the same manner as the proteomic dataset), whereas the plasma samples were annotated according to the corresponding tissue sample. Sparse partial least squares discriminant analysis (sPLS-DA, ^27^) of these plasma samples showed there is good discrimination between the high and low proliferation groups, indicating that tumor proliferation status is reflected in the plasma proteome (**Figure 4D**).

**Figure 4.**
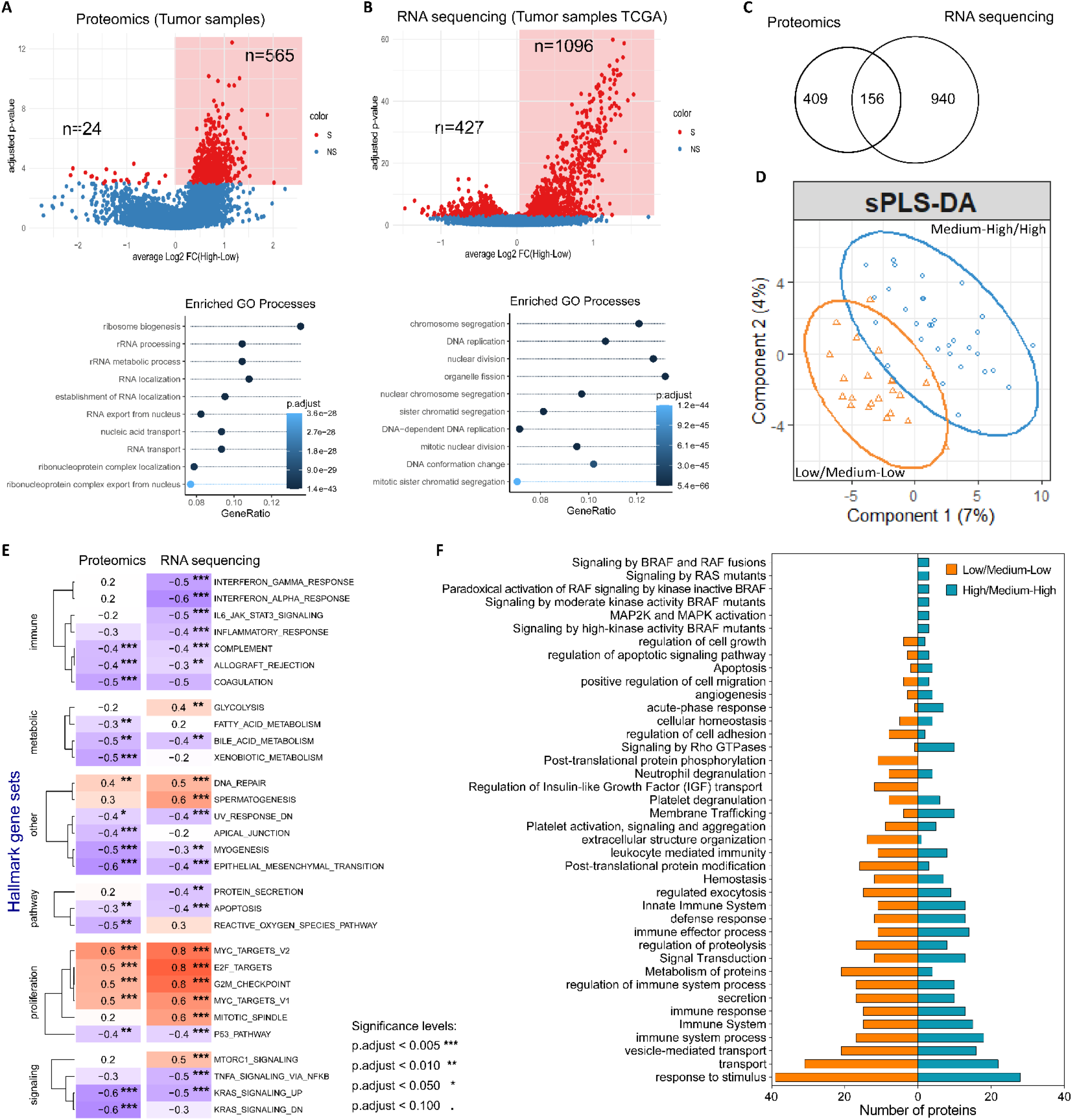
Biological processes altered with increased proliferation are similar across tumor proteome, tumor transcriptome, and plasma proteome. **A)** Volcano plot of the −log10 transformed ANOVA test p-value (FDR-corrected) and the average log_2_ FC_High-Low_ of the proteins, and the Gene Ontology Biological Process (GO BP) over-representation analysis of the significant proteins (FDR < 0.001). **B)** Volcano plot of the −log10 transformed ANOVA test p-value (FDR-corrected) and the log_2_ FC_High-Low_ of transcripts, as well as the GO BP over-representation analysis of the significant transcripts (FDR < 0.001). **C)** Overlap between significantly overexpressed proteins and transcripts in higher proliferation groups. **D)** sPLS-DA on melanoma plasma proteome. Ellipses delimit the sample groups created based on the proliferation status of the corresponding tumor samples. **E)** Enrichment scores for hallmark gene sets as provided by pre-ranked GSEA. The analysis provides further evidence that similar processes are up-and downregulated with increased proliferation in both data sets. **F)** Biological pathways and processes significantly enriched in the top plasma proteins discriminating between high and low proliferation.

Differential expression analysis resulted in 598 proteins (**Figure 4A**) and 1523 transcripts (**Figure 4B**) with significant differential expression across the proliferation groups (ANOVA test FDR < 0.001). The overlap between these significant components was rather sparse on the gene level (**Figure 4C**), however, this list of proteins and genes both suggest an upregulation of cell cycle processes. Also, similar upregulation of proteins that were involved in the organization and localization of cellular components, and an increased biosynthesis of nucleotides. The latter process is particularly important for proliferation because nucleotides cannot be directly taken up from the extracellular space in larger quantities ^28^. We additionally performed pre-ranked Gene Set Enrichment Analysis (GSEA) to see which MSigDB hallmark gene sets are concordantly up-, or down-regulated in highly proliferating tumors. The majority of these processes received similar enrichment scores, in both omics data (**Figure 4E**), strengthening the idea, that RNA-level data from TCGA is a valuable resource for supplementing our protein-level results. We observed a systematic downregulation of immune-related processes, which is associated with increased proliferation, possibly due to a lack of immunoediting at a more progressed stage of the disease ^29^. The relation between immune system modulation and tumorigenesis in melanoma has been studied for the past few years, where today immunotherapies are the most effective treatment options for melanoma patients ^30^. Our observation provides evidence, supported by the high mutational load, that melanoma development and proliferation rely on evasion of the immune surveillance by downregulating immune response-related pathways.

Plasma proteins with altered abundance profiles across proliferation groups were delineated by selecting the top 100 proteins that drive the discrimination of groups according to the sPLS-DA analysis. These proteins are related to the: immune system, extracellular structure organization, vesicle-mediated transport, platelet degranulation and activation, acute-phase, cell migration, and apoptosis (**Figure 4F**). It has been shown that the platelets play a central role in the tumor growth and metastasis development in, melanoma, along with additional other cancer diseases. There is evidence postulating that tumor cells interact with platelets, thereby inducing aggregation as well as degranulation ^31–34^. Regarding the extracellular structure organization, it has been shown that extracellular matrix proteins have a role in the cell transformation process since they are responsible for the interaction between cells and the microenvironment ^35^. Moreover, inflammatory process and cancer development are inter-connected, positioning acute-phase proteins as potential biomarkers in several types of cancer ^36^. Some members of these plasma proteins are prognostic serum/plasma biomarkers (LDH ^37^, e.g., Serum amyloid A ^36,38,39^), with increased expression in patients with short survival (CRP ^36,40^, as well as ORM1 ^41^). Notably, all these proteins were identified as upregulated in the medium-high/high proliferative groups, which indicates a correlation to tumor cell proliferation. The observation that the main plasma proteins that discriminate between groups of tumors based on proliferation are mainly involved in biological pathways and processes related to the response of the immune system, has strong implications on the influence that melanoma exerts on the immune system, outside their physical limits.

### 2.5. Study of the melanoma kinome in the context of tumor proliferation

The study of the kinome through proteomics and phosphoproteomics provided further insight into the interplay between tumor proliferation and the repressed immune system. We first explored the kinome data of our sample cohort (including non-tumor and tumor microenvironment samples) comprehensively. We identified members of each major protein kinase class, with a fairly even distribution (**Figure 5A**). Out of the 522 human kinases present in the KinMap database ^42^, we were able to directly identify 334 kinases, from which 297 and 232 were quantified in our proteomic and phosphoproteomic data respectively. Furthermore, we performed motif analysis using motifeR ^43^ on the phosphorylated peptides and found 104 enriched motifs. These are predicted to be substrates of 260 kinases, out of which 58 kinases were not experimentally identified in our study. The number of identified and predicted kinases thus reached 75% of all human kinases, which indicates that our data of the melanoma proteome and phosphoproteome can capture the majority of phosphorylation events occurring in melanoma (**Figure 5B-C**). The GMGC group is an interesting kinase family where many kinases were detected at the protein level, which we also found to be upregulated in highly proliferating tumors. This group entails cyclin-dependent kinases (CDK) as well as mitogen-activated protein kinases (MAPK), both mediators of cell proliferation ^44,45^.

**Figure 5.**
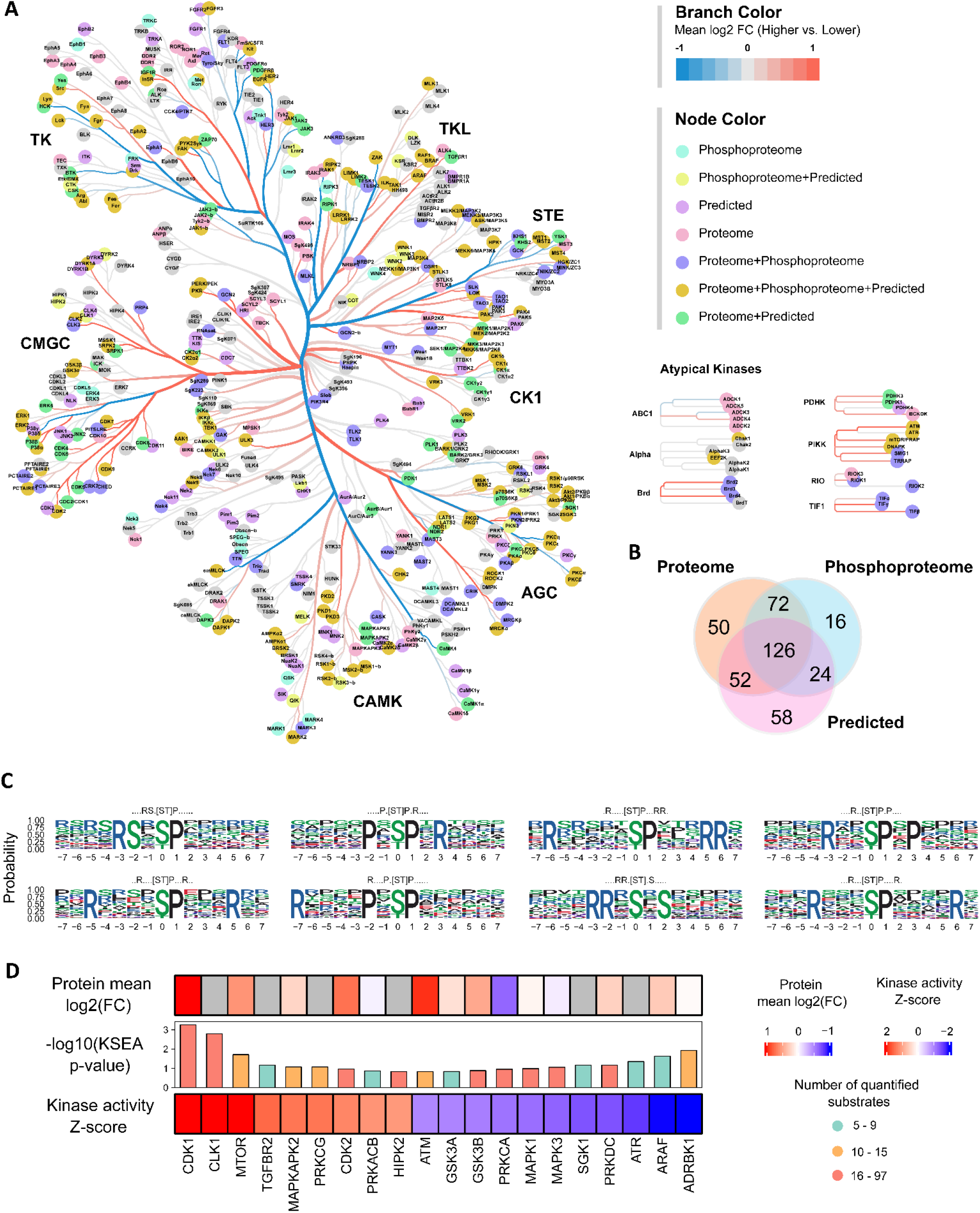
Kinase activities altered with increased proliferation. **A)** A kinome tree illustrating the wide range of kinases identified and predicted in our multi-omic study of tissue samples (including non-tumor and tumor microenvironment samples). Coloring of the branches was according to the mean log2 FC (higher vs lower proliferation status) of the kinases. The figure was made using Coral ^59^. **B)** Venn diagram indicating the number of kinases directly identified in our proteomic and phosphoproteomic data, and the overlap between the predicted and identified kinases. **C)** The top 8 enriched motifs in the phosphoproteome. The peptide sequence motif plots were generated using motifeR. **D)** Inference of relative kinase activities between higher and lower proliferating tumors using kinase-substrate enrichment analysis. Here we show only the kinases for which at least 5 substrates were detected in the phosphoproteomic data and showed significantly altered activity (p-value of the Z-score < 0.15). The total amount of detected substrates (phosphopeptides), as well as the protein abundance change for each kinase between high and low proliferation groups (if available) is also noted. Grey colored cells represent a missing value.

Turning our focus only to the tumor tissue samples, we were interested in the changes of kinase activities between high and low proliferating tumors. We performed kinase-substrate enrichment analysis using the KSEA App ^46–49^ which allows us to computationally infer the relative kinase activity based on the quantitative phosphoproteomic data. We identified 20 kinases for which at least 5 substrates were quantified in our data, and also showed altered activity (the p-value of the kinase Z-score was less than 0.15) (**Figure 5D**). Many of these kinases are well-known to be directly or indirectly regulators of cell proliferation-related processes, as expected. Interestingly, we identified multiple kinases involved in the immune system which has been mentioned in the literature as notable players in immunotherapies. A recent study suggested that CDK1 might contribute to a mechanism that induces immune escape in hepatocellular carcinoma and could be utilized to predict the patient’s response to immunotherapy ^50^. Another study on SKOV3 ovarian carcinoma cells proposed that CDK2 (and CDK4) may play an important role in the proliferation-promoting effects of tumor-associated macrophages ^51^. mTOR is viewed as a potential oncogene in targeted immunotherapies as studies suggest that the mTOR signaling is vital in the functions of the immune cells ^52^. The GSK3 kinase (ergo its two isoforms, GSK3A and GSK3B) have been proposed to be a good candidate for immunotherapy, as its inhibitors promote an immune response against a wide range of human cancer cells, including melanoma ^53^. The RAF kinases (ARAF, BRAF, CRAF) are intensively studied due to their role in tumorigenesis regulation, however, it was also found that they are required for the differentiation and function of dendritic cells ^54^. PRKCA was shown to be involved in the proliferation and differentiation of immune cells, although its function shows both pro-and anti-oncogenic properties ^55^. Moreover, PRKCA phosphorylates TRPC1 and thus regulates the Ca2+ entry in human endothelial cells ^56^. A research study recently suggested that tumors that harbor loss-of-function PRKDC mutations, including melanoma, may benefit from immune therapy ^57^. The inactivation of SGK1 can be associated with anti-oncogenic effects both on tumor cells and on the immune microenvironment and is considered a potential target in non-small cell lung cancer ^58^. In summary, our kinome data suggested the importance of a couple of kinases in melanoma cell proliferation and the regulation of the immune system. To the best of our knowledge, not all kinases were previously connected to melanoma, thus, the role of these kinases at the level of immune-regulation in this particular cancer is yet to be established.

### 2.6. Metabolic reprogramming in melanoma

The metabolic shift from oxidative phosphorylation to glycolytic phenotype has been considered an established common feature for most cancers ^60,61^. Previously, it was reported that melanoma metastases upregulate glycolysis compared to primary tumors ^10^. Our observations confirm that melanoma shows upregulation of the glycolysis, moreover, its upregulation correlates with high proliferation (**Figure 6A-B**). The isoforms M1 and M2 of the pyruvate kinase (PKM) gene are produced by differential splicing. PKM-M1 is the main form in muscle, heart, and brain, and PKM-M2 is found in early fetal tissues as well as in most cancer cells. In melanoma we found PKM-M2 upregulated in all tumor sample groups and the tumor microenvironment, confirming the hypothesis that metabolic reprogramming in melanoma extends to the tumor microenvironment (**Figure 6C**). The upregulation of the hexokinases (HK1-3), and the Phosphoenolpyruvate carboxykinase (PCK2) support the accumulation of intermediates of glycolysis required for cell proliferation. However, the enzymes involved in the conversion of pyruvate to Acetyl-CoA were upregulated in tumors, which indicates that part of glycolysis intermediates are directed to the TCA cycle. In addition to the glycolysis, the tumors and their microenvironment significantly upregulated the oxidative phosphorylation (OXPHOS) and the mitochondrial translation (**Figure 6D**). These findings suggest that to support their high proliferation rate, tumors rely on glycolysis and OXPHOS for the production of energy and proliferation-related intermediates. In agreement with these results, we found that most of the protein elements of the hypoxia-inducible factor 1 (HIF-1) signaling pathway were upregulated in tumors (**Figure 7A**). Consequently, the activation of the HIF-1 signaling pathway provides melanoma with a continuously proliferating activation signal, even under conditions of nutrient deprivation.

**Figure 6.**
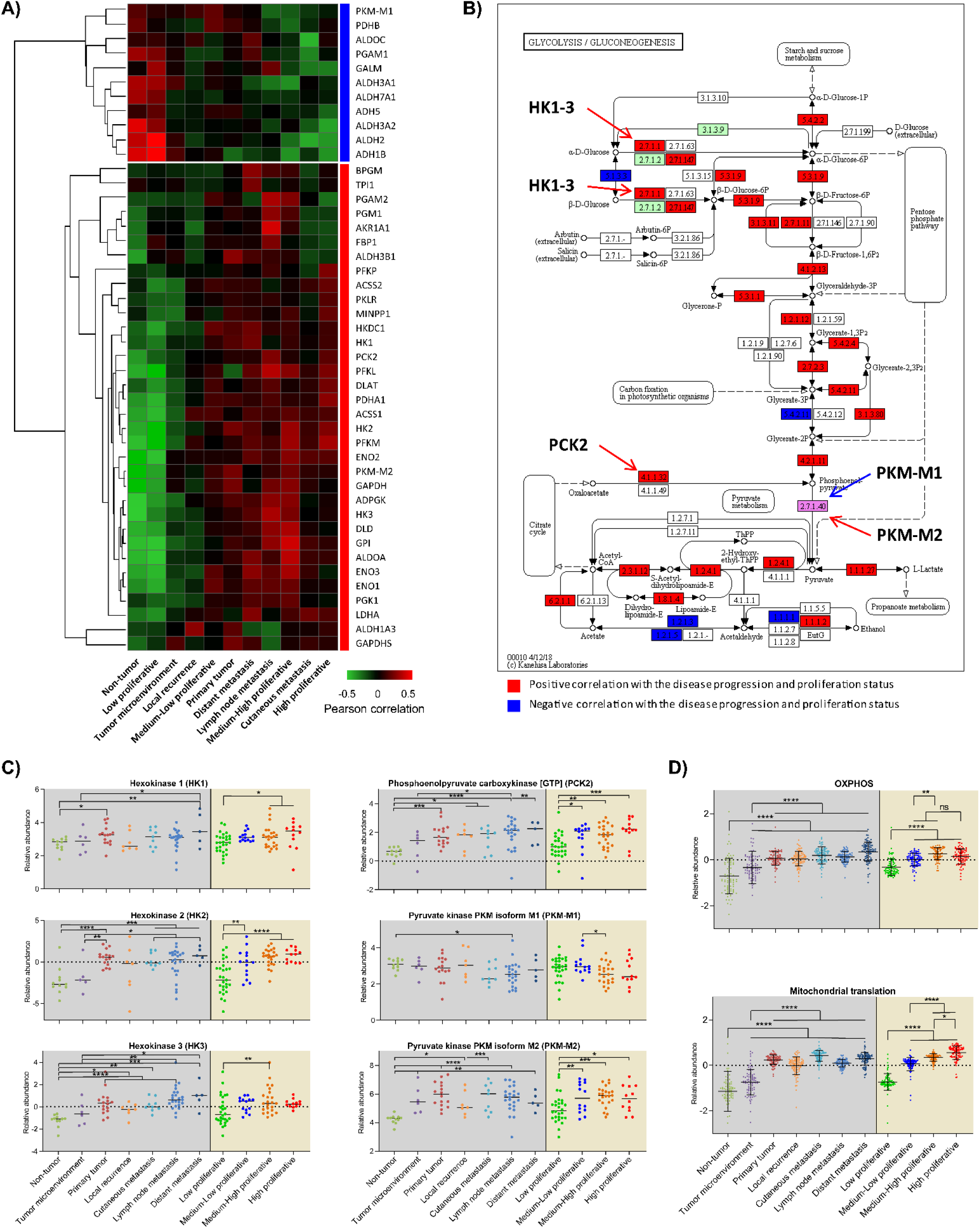
Melanoma progression is characterized by alteration of energy production pathways. **A)** Hierarchical clustering and heatmap of the Pearson correlation values between the relative abundance of proteins reported as part of the glycolysis pathway and the groups of samples based on the sample origin and the proliferation status. Two major protein clusters were identified, those that show a negative correlation (blue rectangle) and those that show a positive correlation (red rectangle) with the melanoma progression and higher proliferation. **B)** Schematic representation of the glycolysis pathway where the proteins identified and quantified in the study were highlighted. The proteins that are marked with red and blue color show a correlation between their abundance profile and the progression of the disease (red: positive, blue: negative correlation) The protein pyruvate kinase PKM is also highlighted in pink, for which two isoforms were identified and PKM-M2 showed a strong positive correlation with the disease progression while the PKM-M1 isoform showed a weak negative correlation. **C)** Relative abundance of proteins across different sample groups. Grouping was both based on the sample origin (from Non-tumor to Distant metastasis) and based on the proliferation status (from Low to High proliferative). Selected proteins were: hexokinases 1-3 (left panels), phosphoenolpyruvate carboxykinase [GTP], mitochondrial, and isoforms M1 and M2 of the pyruvate kinase PKM (right panels). **D)** Mitochondrial oxidative phosphorylation and translation machinery pathways altered across different sample origins and proliferation groups. Proteins reported to be involved in the Oxidative phosphorylation pathway (upper panel) and those involved in mitochondrial translation (bottom panel) were plotted with their mean relative level in the groups of samples. *, **, *** and **** means significant differences between groups (p<0.05, p<0.01, p<0.001 and p<0.0001 respectively).

**Figure 7.**
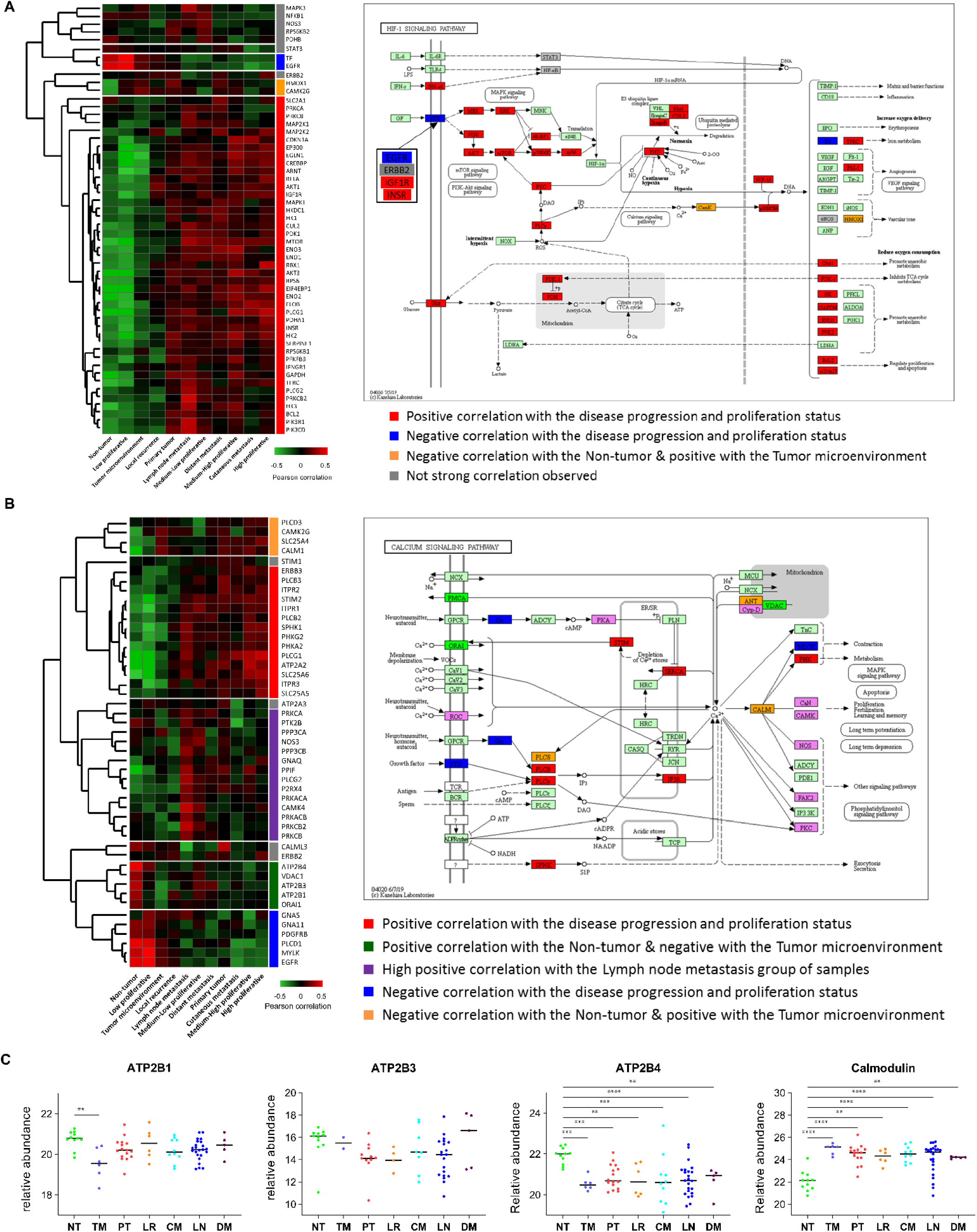
Hypoxia-inducible factor 1 and Calcium signaling pathways are dysregulated during melanoma progression. **A)** Schematic representation of Hypoxia-inducible factor 1 signaling pathway where proteins identified in the study are highlighted. Color indicates the correlation observed between the protein levels and the groups of samples (see legend). **B)** Schematic representation of Calcium signaling pathway where proteins identified in the study are highlighted. Color indicates the correlation observed between the protein levels and the groups of samples (see legend). **C)** Relative levels of the ATPases 1, 3 and 4 (ATP2B1, ATP2B3 and ATP2B4) and calmodulin in groups of samples based on the origin.

Moreover, the calcium signaling pathway which impacts the metabolism and the proliferation of the cell was found dysregulated in melanoma (**Figure 7B**). Proteins involved in controlling the levels of intracellular calcium showed dysregulation. Plasma membrane calcium-transporting ATPases 1, 3, and 4 (ATP2B1, ATP2B3, and ATP2B4), involved in the active release of intracellular calcium through the plasma membrane, were downregulated in the microenvironment. In the case of ATP2B4, it was also significantly downregulated in tumor groups compared to non-tumors. On the other hand, the main intracellular calcium-binding protein, calmodulin, was upregulated in the microenvironment and all tumor sample groups compared to non-tumors (**Figure 7C**). These findings highlight the dependence of melanoma on intracellular calcium, at the level of preventing the release of calcium not only from tumor cells but also from their microenvironment.

### 2.7. Lysine acetylation in melanoma

The acetylation workflow allows not only the identification of site-specific lysine acetylation, but also of the stoichiometry impacting the melanoma staging. On average, more than 1100 peptides and 800 proteins were identified carrying acetylation marks in tumor samples, while for non-tumor and microenvironment samples the acetylated peptides and proteins were about 500 and 350 respectively (**Figure 8A**). The abundance distribution of acetylated proteins highlights the fact that with the current technology and workflow, we can detect acetylation sites and quantify their occupancy in relatively higher abundant proteins compared to the total of identified proteins (**Figure 8B**). This is a reflection of what has been previously reported, that acetylation in lysine residues is a post-translational-modification that generally shows very low occupancy on their target sites ^13,62,63^. In all the samples submitted to the analysis the site-specific occupancy distribution was very similar. At least 50 % of the acetylated peptides identified in all samples, showed a stoichiometry of less than 15% (**Figure S5**), which is also highlighted after normalization of the data using probit transformation ^64,65^ (**Figure 8C**).

**Figure 8.**
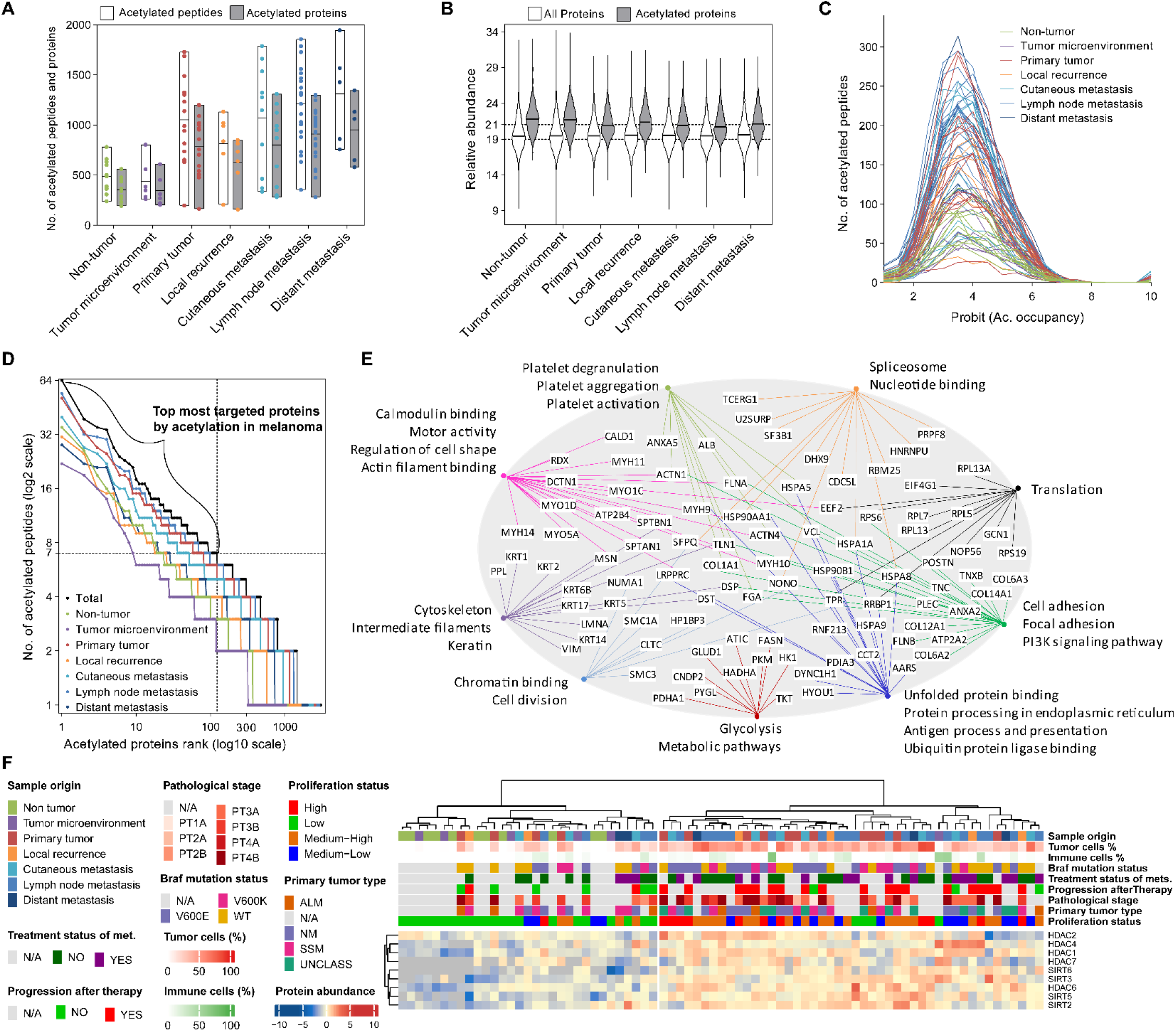
Lysine acetylome in melanoma. **A)** Identified acetylated peptides and proteins in melanoma-related tissue biopsies, distributed based on sample origin. **B)** Abundance distribution based on the sample origin of all identified proteins and those detected being acetylated. **C)** Frequency distribution of site-specific occupancy of acetylated peptides in each sample. The original values in percentage of the site modified were probit transform before plotting to fit normal distributions. **D)** Distribution of acetylated proteins based on the number of acetylated peptides identified in melanoma-related biopsies. Dashed line delimits the proteins with at least 7 different acetylated peptides in the study (122). **E)** Most targeted proteins with acetylation as a post-translation modification in melanoma are connected to their most relevant enriched functional annotation clusters. The enrichment analysis was performed in DAVID Bioinformatics ^74^. **F)** Hierarchical clustering of the tissue samples based on the relative protein abundance of the nine enzymes with reported deacetylase activity, identified in at least two-thirds of the samples.

The melanoma acetylome reported here includes about 3000 proteins, in most of them, only one acetylation site was detected. However, it was found that several proteins have multiple acetylation sites (**Figure 8D**). Particularly, the 100 topmost acetylated proteins were found with at least 7 different acetylated peptides. A functional annotation enrichment analysis of the proteins with at least 7 acetylation sites in melanoma reveals that acetylation regulates pathways involved in the cytoskeleton, its organization, the actin filaments, and the regulation of cell shape. Besides, as previously reported, a large number of proteins from the translation machinery, the spliceosome, and metabolic enzymes were found to have multiple acetylation target sites ^66,67^ (**Figure 8E**). Among the proteins with more acetylation sites in melanoma, we also found a cluster of biological annotations that includes the protein processing in the endoplasmic reticulum, and the antigen process and presentation to be significantly enriched. Particularly these pathways are among the most mutated and dysregulated in melanoma. These findings reveal an important role of lysine acetylation in the regulation of critical pathways for melanoma development and progression.

From the proteomic data, we were able to quantify in at least two-thirds of the samples 9 out of the 18 deacetylases reported in the human genome. HDAC1 and 2 from class I, HDAC4 and 7 from class IIa, HDAC6 from class IIb, and four out of the seven sirtuins or class III HDACs (SIRT2, 3, 5, and 6). The role in cancer of several members of these families has been highlighted ^68–70^. In melanoma, several clinical studies have shown benefit to the patients by targeting HDAC class I, particularly for those with acquired resistance to targeted therapy ^71^. The combination of HDAC inhibitors with immunotherapy has shown an enhanced response to the therapy ^72,73^. Here we show that most of these enzymes, not only are upregulated in melanoma but also their abundance is higher in the more proliferative groups of tumors (**Figure 8F**). The relative level of the deacetylase enzymes in melanoma together with quantitative analysis on the site-specific acetylation targets allow a better understanding of how the treatment with HDAC or specific target inhibitors can be implemented in a personalized treatment strategy.

### 2.8. The melanoma progression involves repression of the tumor amino acid metabolism pathway and the evasion of the immune surveillance

Primary tumors, when compared to non-tumors upregulate proliferation-related pathways such as DNA replication, ribosome biogenesis, spliceosome, and RNA transport. Oppositely, tumors downregulate a set of proteins involved in the communication with the extracellular environment and cell adhesion (**Figure 9C**). The changes in the metabolism of the primary tumors include downregulation of pathways linked to the degradation of amino acids and fatty acids. When compared to non-tumors, although the tumor microenvironment samples do not show upregulation of proliferation-related pathways as primary tumors do, they show similar alterations in the pathways related to the extracellular matrix communication (**Figure 9A-C**). Interestingly, similar pathways and biological annotations of the proteins were significantly enriched in proteins dysregulated at the level of acetylation in their lysine residues. The glycolysis pathway, which upregulation has correlated with the development of melanoma, was found under-acetylated in both primary tumors and their microenvironment compared to non-tumors. Similar results were obtained for ribosomal and translational proteins, suggesting that acetylation on those proteins could primarily have a downregulatory effect on the pathways. Oppositely, several proteins involved in the extracellular matrix and cell-cell adhesion, downregulated in primary tumors showed increased levels of acetylation (**Figure 9D-F**).

**Figure 9.**
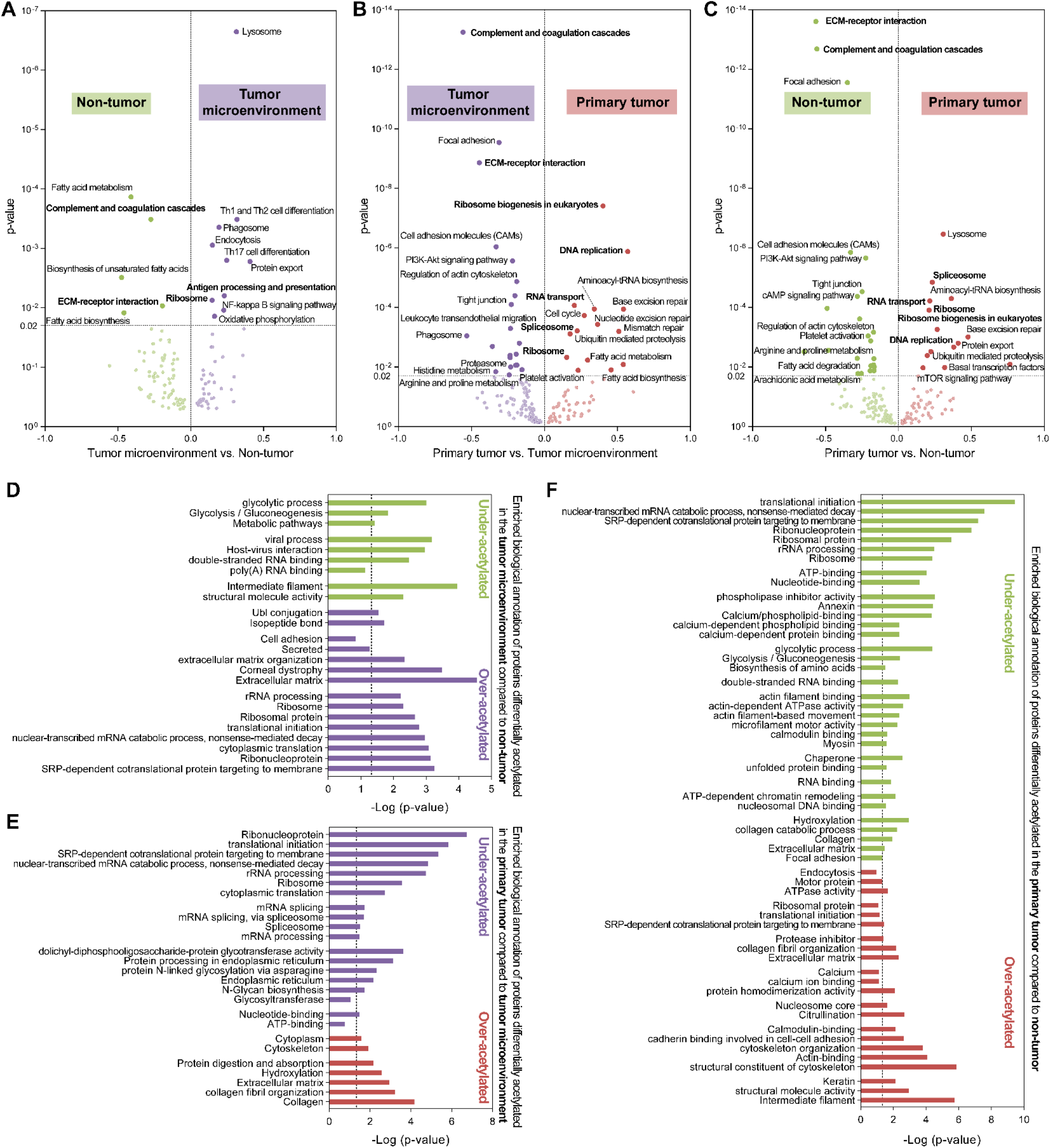
Dysregulated proteome and acetylome in primary melanoma and their microenvironment. Biological pathways significantly enriched when comparing the mean protein abundances between the group of samples based on their origin, according to 1D functional enrichment analysis provided by Perseus software ^75^. Data is represented as volcano plots where the annotation enrichment p-value and the score of the differences were plotted. A false discovery rate (FDR) of 0.02 was set as the cut-off for significance. **A)** Comparison between the tumor microenvironment and the non-tumor samples, **B)** between the primary tumor and the tumor microenvironment, and **C)** the primary tumor and the non-tumor. **D-F)** Functional annotation clusters significantly enriched in acetylated proteins that showed significant differences in their site-specific acetylation occupancy between origin-based groups. D) Comparison between the tumor microenvironment and the non-tumor samples, E) between the primary tumor and the tumor microenvironment, and F) the primary tumor and the non-tumor.

Among the most mutated genes in melanoma identified as hot spot, it was highlighted several members of the antigen presenting MHC class I and class II. A quantitative analysis from the proteomic data reveals that when taken together, the members of the MHC class I on average show lower level in the metastasis groups compare to both non-tumor and primary tumors, although not all the comparisons were statistically significant (**Figure 10A** (left panel)). Moreover, when analyzing only the tumor samples grouped based their proliferation groups, there is a downregulation of this proteins in relation to the proliferation. The average repression of this proteins, between the high and low proliferative groups, was found to be significant (**Figure 10A** (right panel)). Similar results were obtained from the analysis on the identified family members of the MHC class II (**Figure 10B**). These findings provide more insight into how melanoma manages to evade the immune surveillance, by heavily mutating and downregulating the antigen processing and presenting machinery, which ultimately affect the immune system response ^76^.

**Figure 10.**
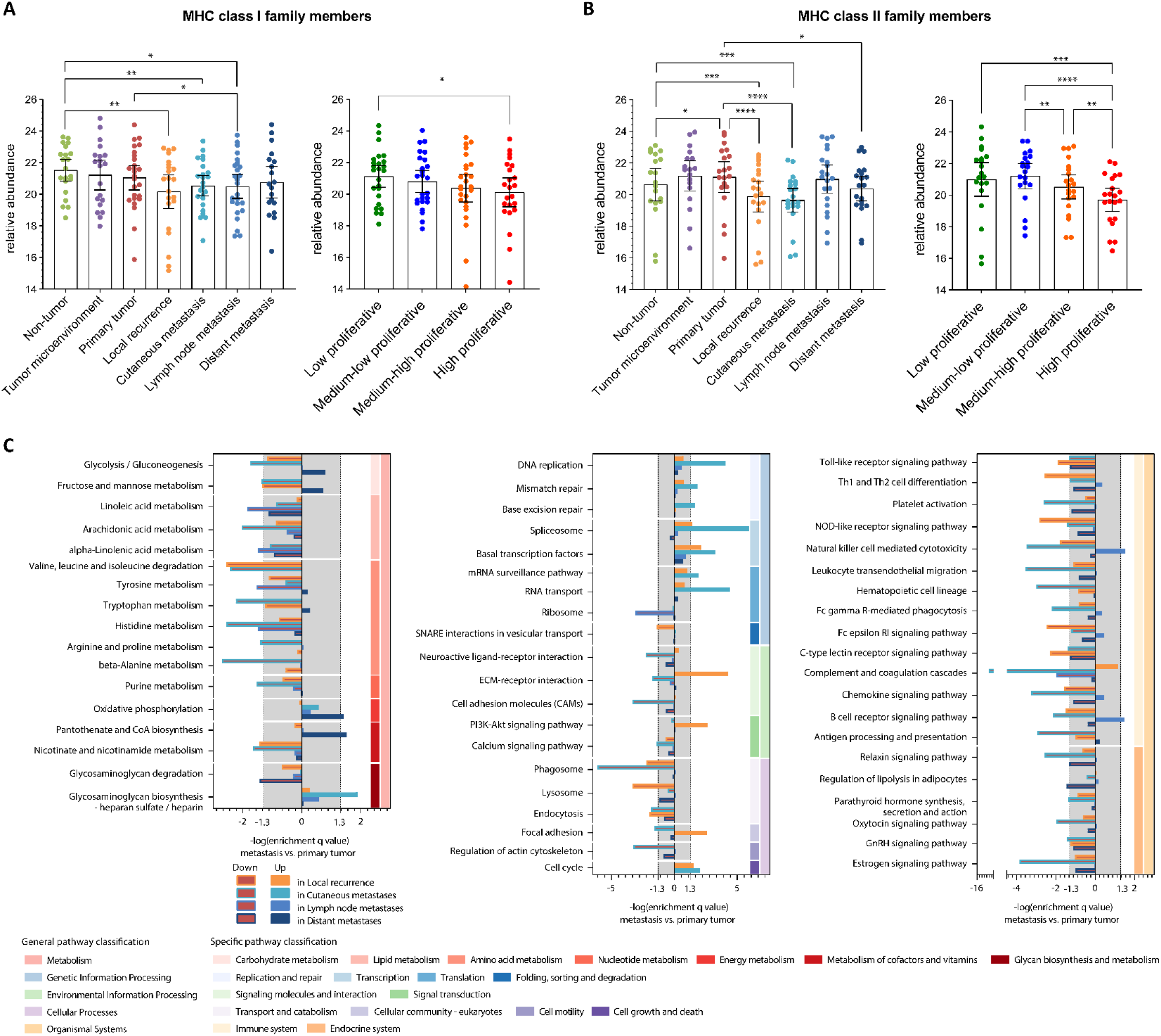
Melanoma progression toward metastasis is driven by changes in the metabolism and downregulation of the immune system-related pathways. Relative levels of the identified family members of the MHC class I **(A)** and class II **(B)**. Samples were grouped based on their tissue origin (left panels), and based on their proliferation status (right panels). **C)** Dysregulated biological KEGG pathways between the primary tumors and the different groups of metastases included in the study. The pathways were distributed according to their classification (general and specific). Bar length correspond to the –log10 (q-value), and the gray area correspond to non-significantly enriched. Negative and positive values correspond to down-and upregulated in the metastasis compared to the primary tumor group, respectively.

The progression of melanoma from primary lesions towards metastasis was studied through a biological pathway enrichment analysis on dysregulated proteins between metastases and primary melanomas. Represented pathways were found significantly dysregulated in at least one of the metastasis groups compared to primary tumors (**Figure 10C**). The metabolic changes in metastatic melanoma continue with the downregulation of the amino acid metabolism, as observed in the comparison between the primary and non-tumor samples. Similarly, pathways linked to the communication with the extracellular environment, including proteins involved in cell-cell adhesion were downregulated in metastases. In this sense, the repression of amino acid metabolic and cell communication pathways is a common feature of melanoma development and progression. Additionally, one of the most relevant characteristics of metastatic melanomas compared to primary tumors is the downregulation of immune system response-related pathways (**Figure 10C**). The repression of pathways linked to the immune system is a common feature of melanoma proliferation and progression. Pathways involved in DNA replication and repair, transcription, and translation showed upregulation in the metastasis groups, mostly in cutaneous metastases. Altogether, the aggressiveness of melanoma is supported by a combined upregulation of proliferation-related pathways, rewiring of the metabolism, and repressing the immune system response.

## 3. Discussion

In this study, we provided new insights into the progression of melanoma through a comprehensive proteogenomic analysis on primary-, and metastasis tumors, including 77 tissue-, and 56 plasma-samples from 47 melanoma patients and with various sample origins (non-tumor, tumor microenvironment, primary tumor, local recurrence, cutaneous, lymph node, and distant metastasis). We included deep proteomics and PTM data accompanied by whole exome sequencing to provide an integrated characterization of MM. Several studies have been conducted in large melanoma cohorts using high throughput omic analyses ^8,9,77^ to gain a better understanding of the molecular mechanism that underlies the development and progression of melanoma. We emphasize the processes that are dysregulated with increased melanoma cell proliferation. Hence, we inferred the proliferation status of each tissue sample based on the relative protein levels of the MCM complex elements. The plasma samples originating from patients with high or low proliferation tumors were also distinguishable at the plasma proteome level. Including the Skin Cutaneous Melanoma TCGA PanCancer data in the analysis, we further validated and supplemented our findings using a larger sample cohort.

Whole exome sequencing of the tumor samples highlighted that mutations occurring in melanoma stem from similar underlying molecular mechanisms in all samples. Hot spot analysis identified 132 genes with frequent somatic mutations, and notably, many members of this list were known to be interactors or regulators of the immune system, such as the MHC class I and II elements. Furthermore, the signaling cascades (MAP kinases and PI2K-AKT) and the calcium homeostasis were the most heavily dysregulated pathways due to the high mutation burden in melanoma. These observations on exome level were reaffirmed on protein level. Firstly, we systematically identified proteins and genes that indicate the suppression of the immune response throughout our analyses, both during the exploratory analysis of the tissue and plasma proteome and the TCGA data, and both during the more targeted analyses. Our findings point to a direct link between the repression of members of the MHC family by mutations or translational downregulation, and progression of melanoma. Moreover, we found this repression in correlation to tumor proliferation. The relation between immune system modulation and tumorigenesis in melanoma has been extensively studied for the past few years, as increased proliferation and lack of immunoediting go hand in hand at a more progressed stage of the disease ^78–80^. While immunotherapies are the most effective treatments for melanoma patients nowadays, far from all patients respond favorably, and there is therefore still a need for increasing our knowledge of melanoma biology to develop new therapies, strategies to overcome resistance, and increase the number of patients responding to the immunotherapeutic agents ^29,30^.

The quantitative analysis of two of the most important posttranslational modifications, the phosphorylation and lysine acetylation, deeply complements the proteomics analysis by adding elements of regulation at a level usually ignored by other large-scale analyses. Our analysis of the melanoma kinome highlighted that 75% of all human kinases were detected or predicted based on their phosphorylated targets, which allows to capture the major phosphorylation events. The kinase-substrate analysis outlined multiple kinases that are activated or deactivated with increased proliferation. Moreover, several of these kinases are considered potential therapeutic targets due to their role in the function of the immune system, such as the GSK3A kinases or the SGK1. Additionally, to the best of our knowledge, this is the first lysine acetylome stoichiometry analysis reported for a large number of melanoma tumor samples. Our observations highlight that the translation machinery and metabolic enzymes are regulated by acetylation and as a general trend lower acetylation degree resulted in upregulation of the pathway. This is particularly interesting because several of the deacetylase enzymes identified in the study were indeed upregulated in melanoma. Our findings support previous reports that HDAC inhibitors treatment in melanoma particularly in combination with immunotherapy enhance the response of the patients ^72,73,81^.

We found that both glycolysis and OXPHOS are utilized by highly proliferating melanoma cells to produce energy and the proliferation-related intermediates, supported by the observed activation of the HIF-1 signaling pathway, leading to continued proliferation even in the event of nutrient deficit. We further characterized melanoma by the downregulation of proteins involved controlling the levels of intracellular calcium, the repression of amino acid metabolism, and the suppression of cell communication. The observed metabolic reprogramming extended to the tumor microenvironment, but was not discovered in non-tumor samples.

We acknowledge the limitations of our study. An experimental design involving samples from 7 different tissue origins is ultimately more prone to bias due to the low sample sizes within each sub-cohort. Nonetheless, the integration of genetic level information and PTM data next to proteomics represents a rich resource to investigate melanoma biology, and main observations from one type of omic data were supplemented on another molecular layer, thus confirming the validity of our results. We envision that the hypotheses underlined in this manuscript will contribute to a better understanding of melanoma and will aid the development of new therapies.

## 4. Experimental section

### 4.1. Sample processing and peptide/protein identification

A protein extraction buffer containing SDS 2%, DTT 50 mM, Tris 100 mM, pH 8.6, was added to the sliced tissues or cell pellet, rest for one minute, and sonicated using a Bioruptor plus (40 cycles, 15 s On, 15 s Off, at 4 °C). Samples were incubated at 95 °C for 5 min. Proteins were alkylated by adding IAA to a final concentration of 100 mM for 20 min in the dark at room temperature (RT). The proteins were precipitated overnight by adding 9 volumes of cold ethanol. Proteins were dissolved in TEAB buffer containing SDS 1%, SDC 0.5%, and submitted to a chemical acetylation reaction of all free amino groups. The reaction was performed with N-acetoxy-succinimide-d3 (NAS-d3). O-acetylation was reverted by treating the samples with 5% hydroxylamine. Proteins were precipitated as described above to remove the excess reagents and dissolved in AmBiC containing SDC 0.5%. Proteins were digested with trypsin (1:50, enzyme:protein) at 37 °C, overnight. The SDC was removed from the peptide solution by adding ethyl-acetate and TFA ^13^. After discarding the organic phase, the mixtures of peptides were quantified and kept at −80 °C until MS analysis.

For phosphopeptide enrichment, 80 mg of peptides was submitted to an automated workflow, performed on an AssayMAP Bravo system (Agilent Technologies). The procedure was performed as previously described ^11^. Phosphopeptides were dried and kept at −80 °C until MS analysis.

#### Peptide spectral library

Pools of peptides including all the tumor samples were spiked with iRT and analyzed in a high-resolution LC-MS system (Dionex Ultimate 3000 RSLCnano UPLC coupled to a Q-Exactive HF-X mass spectrometer (Thermo Fischer Scientific)). Peptides were desalted on a trap column Acclaim PepMap100 C18 (3 µm, 100 Å, 75 µm i.d. × 2 cm, nanoViper) and then connected in line with an analytical column EASY-spray RSLC C18 (2 µm, 100 Å, 75 µm i.d. × 50 cm). The temperature of the trap column and analytical column was set at 35 °C and 60 °C respectively, and the flow rate was 300 nL/min. A non-linear gradient of 137 min was used for peptide elution. Data were acquired applying a DDA method covering a mass range of 385-1460 m/z. The parameters for MS1 included a resolution of 120,000 (@ 200 m/z), the target AGC value of 3·10^6^, and the maximum injection time was set at 100 ms. For MS2, the resolution was 15,000, the AGC value 1·10^5^, the injection time 50 ms, the threshold for ion selection 8·10^3^, and the normalized collision energy (NCE) 28. The isolation window was 1.2 Th.

#### DIA analysis

samples were analyzed in duplicates using the same LC-MS system and with the same chromatographic conditions as described above. Data were acquired using a DIA method intended for peptide quantification using MS1 scans. The DIA method contains 54 variable width MS2 windows, based on the empirical distribution of peptides, and 3 full scan MS1. A full MS1 scan was placed every 18 MS2 scans. The parameters for MS1 were: resolution 120,000 (@200 m/z), AGC target 3E+06, injection time 50 ms, range 385-1460 Th. For the MS2 were: resolution 30,000, AGC target 1·10^6^, injection time 50 ms, NCE 28.

Protein and peptide identification and quantification were performed with the Spectronaut software (Biognosys, AG). DDA raw files were searched against a human protein database downloaded from UniProt in 2018, to create a spectral library. The search engine used was pulsar, which is integrated into the Spectronaut platform. The parameters included Arg-C as the cleavage enzyme, 2 missed cleavages were allowed, lysine acetylation (d3) and carbamidomethyl-cysteine were set as fix modifications, methionine oxidation and protein N-terminal acetylation (d3) and (d0) were set as variable modifications. For the postmortem cohort, the selected enzyme was trypsin, two missed cleavages were allowed, carbamidomethyl-cysteine was fixed, and methionine oxidation and protein N-terminal acetylation were set as a variable. The identifications were controlled by an FDR of 1% at the PSM, peptide, and protein levels.

For identification and quantification of lysine acetylation, raw files were analyzed by two different software Pview ^82^ and Proteome discoverer (Thermo scientific). Acetylation stoichiometry was only considered for those peptides commonly identified by the two search engines. The parameters included Arg-C as the cleavage enzyme, carbamidomethyl cysteine as fix modification, methionine oxidation, and lysine acetylation (d0 and d3) were set as variable modifications. The number of identifications was controlled by FDR of 1% at peptide and protein level. The parameters for the stoichiometry calculation were as previously reported ^13^. For comparison of the acetylation occupancy between groups, a probit transformation of the data was applied.

### 4.2. Proteomic, phosphoproteomic, and transcriptomic data processing, statistics, and bioinformatics

The data processing and statistics were performed in R (vs. 4.0) unless specified otherwise. The Skin Cutaneous Melanoma TCGA PanCancer data ^9,83,84^) was downloaded from the cBioPortal website ^85,86^.

A similar data pre-processing workflow was applied on protein and phosphopeptide intensities as well as on the transcriptome data. For the RNA expressions, counts equal to 0 were treated as NA values. Intensities/counts were log_2_ transformed, followed by global median centering for normalization, and sample replicates averaging in case of the proteomic data. The correlation in abundance between proteins from the MCM protein complex was used as a criterion to evaluate the normalization performance. Differential expression analysis was performed using Analysis of Variance (ANOVA test), followed by ANOVA p-value adjustment with the Benjamini-Hochberg method and an additional Tukey-Kramer post hoc test for each ANOVA analysis. No prior filter for valid values was applied on either data set, however, only those groups were compared where the protein/phosphopeptide/gene had valid values in at least 70% of the groups’ samples. Comparison between proteomic and transcriptomic signatures related to proliferation status was done using Gene Set Enrichment Analysis (GSEA) on the 50 hallmark gene sets that were downloaded from http://www.gsea-msigdb.org/gsea/msigdb/index.jsp. The ordered list of mean log2 FC values across the proliferation groups was used as the input and no p-value threshold was set. The hallmark gene sets were further categorized according to Liberzon et al. ^87^. Gene Ontology over-representation test on the biological processes and Gene Set Enrichment Analysis was performed using the R package “clusterProfiler” (vs. 3.18.1).

#### Kinome analysis

The list of human kinases was downloaded from http://www.kinhub.org/kinases.html (on 10 March 2021). Sequence pre-alignment and motif analyses were done using the motifeR web server (https://www.omicsolution.org/wukong/motifeR/). In this analysis, we included only the phosphopeptides that were identified in at least 30% of the samples and where the phosphorylation site was confidently identified (8994 sequences). Sequence data were aligned to have 15-amino-acid long sequences where the central residue is the phosphorylated S/T amino acid. For the motif analysis, the following settings were applied: each motif had to appear at least 20 times in the data set, and the p-value threshold for the binomial probability was set to 1e-06 for the selection of significant residues in the motif. The default human database was used as a background and classical multiple sites analysis was chosen. Motif enrichment analysis was done both on peptides with one and with multiple modification sites. Kinases were predicted based on all the enriched motifs and the minimum NetworKin score was set to 3.

Kinase-substrate enrichment analysis was performed using the KSEAapp R package. Both curated annotations from PhosphoSitePlus and kinase-substrate predictions from NetworKIN were considered (Kinase-Substrate dataset downloaded from the KSEAapp Github page https://github.com/casecpb/KSEA), and the minimum NetworKIN score was set to 5.

### 4.3. WES analysis

*WES data analysis:* The WES sequencing data of the 60 samples were mapped against GRCh38 and analyzed according to standard workflows as described by GATK (Broad Institute). Additional filtering was applied to remove variants with coverage of less than 3 reads. Next, the exome panel enrichment regions from Twist Bioscience (downloaded from https://www.twistbioscience.com/sites/default/files/resources/2019-06/Twist_Exome_Target_hg38.bed) were used to filter out all identified variants outside of these WES targeted regions. The non GCTA nonTCGA_gnomAD subset was used as “population background” information to distinguish between variants annotated in this database referred to as germline and those missing (referred to as somatic variants). Summary statistics about the variants in each of these 2 categories were created using the “bcftools stats” utility.

#### Hot spot analysis

The somatic and germline variant counts were grouped and summed by the gene they overlap with. Chi-square tests were performed to determine genes within individual samples for which the number of somatic variants was different than expected based on the sample’s ratio of these counts across all genes. Similarly, the variant counts per gene were grouped and summed on a pathway level. Bonferroni correction was applied on the obtained p-values. Genes with significant enrichment were selected by using the significance level 0.05 and requiring the presence of significant enrichment in ⅓ of the data (20 out of 60 samples). Data visualization and final statistics were made with a custom R script.

## 5. Acknowledgments

This study was kindly supported by the Berta Kamprad Foundation, Thermo Fischer Scientific as our strategic partner, The Swedish Pharmaceutical Society and Agilent Technologies. This work was done under the auspices of a Memorandum of Understanding between the European Cancer Moonshot Center in Lund and the U.S. National Cancer Institute’s International Cancer Proteogenome Consortium (ICPC)

## 7. Supplemental figures

**Figure S1.**
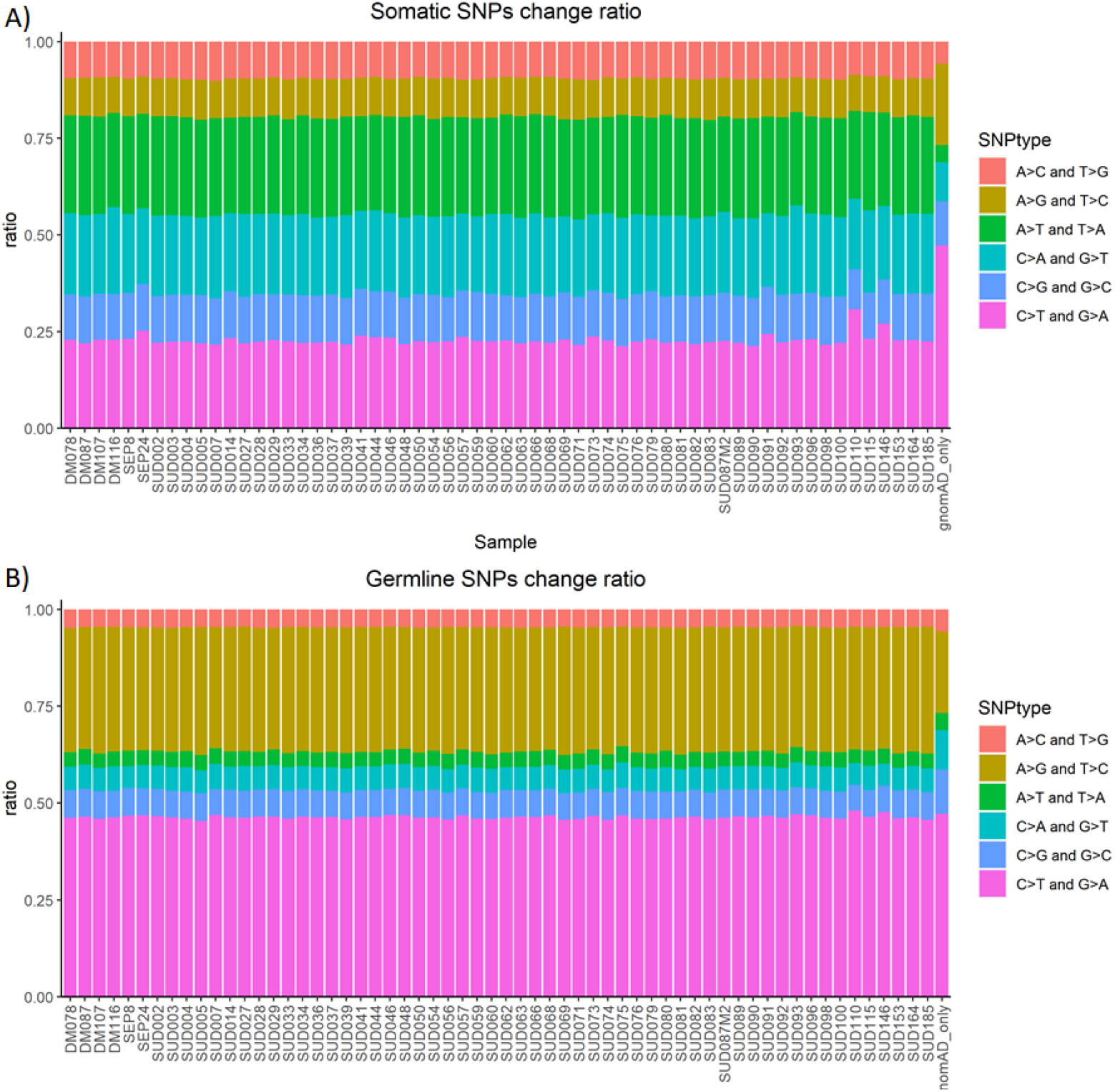
The ratio of various base-pair changes in somatic (A) and germline (B) SNPs compared to population reference (gnomAD_only excluding mutations identified in all tumor samples). The pattern is highly reproducible for both cases with difference between somatic and germline SNPs (A and B).

**Figure S2.**
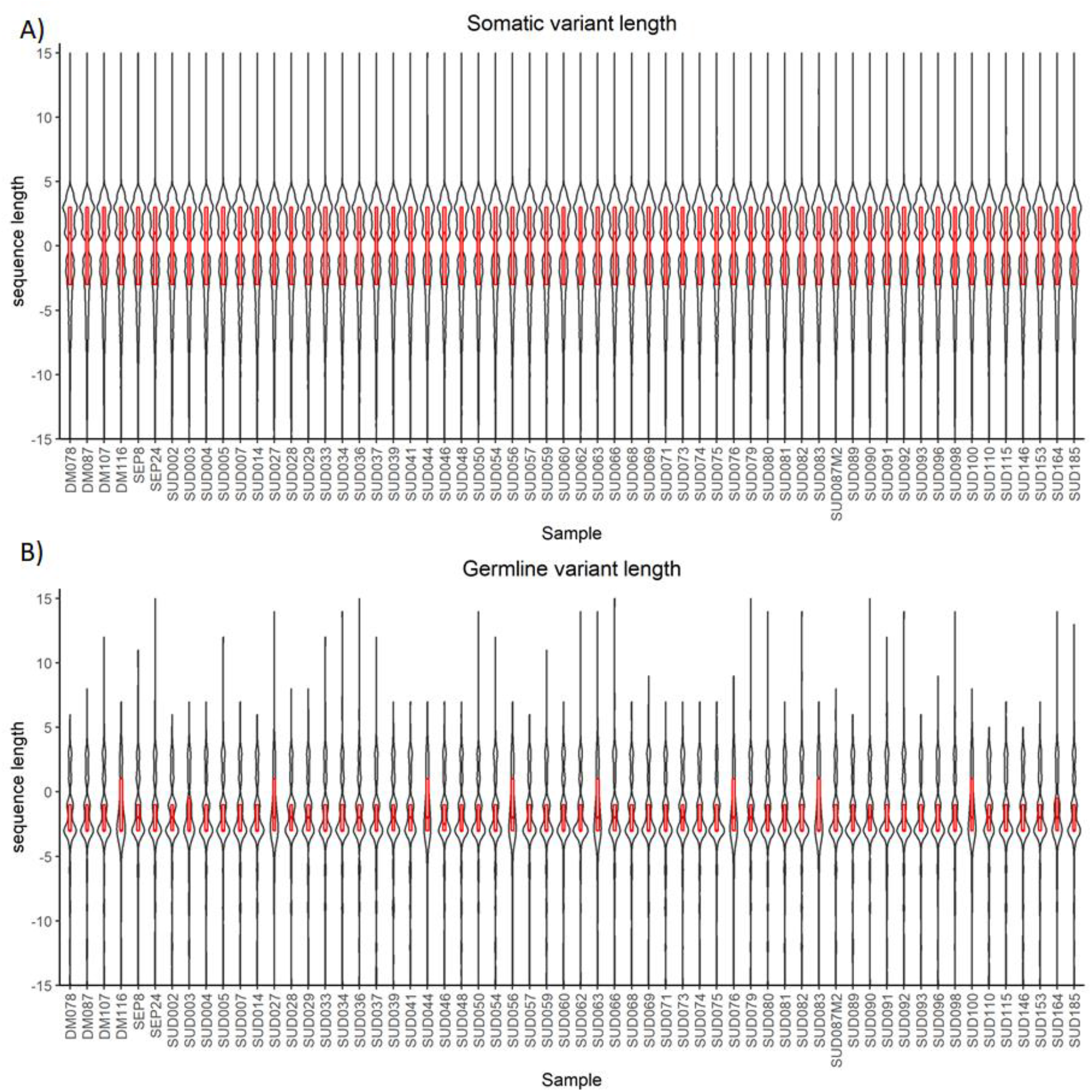
Distribution of sequence length shortening or extension of indels with somatic (A) and germline (B) origin. Somatic indels tend to extend, while germline indels generally shorten exons and protein variants. Y axis show the range of ±15 base pair lengths.

**Figure S3.**
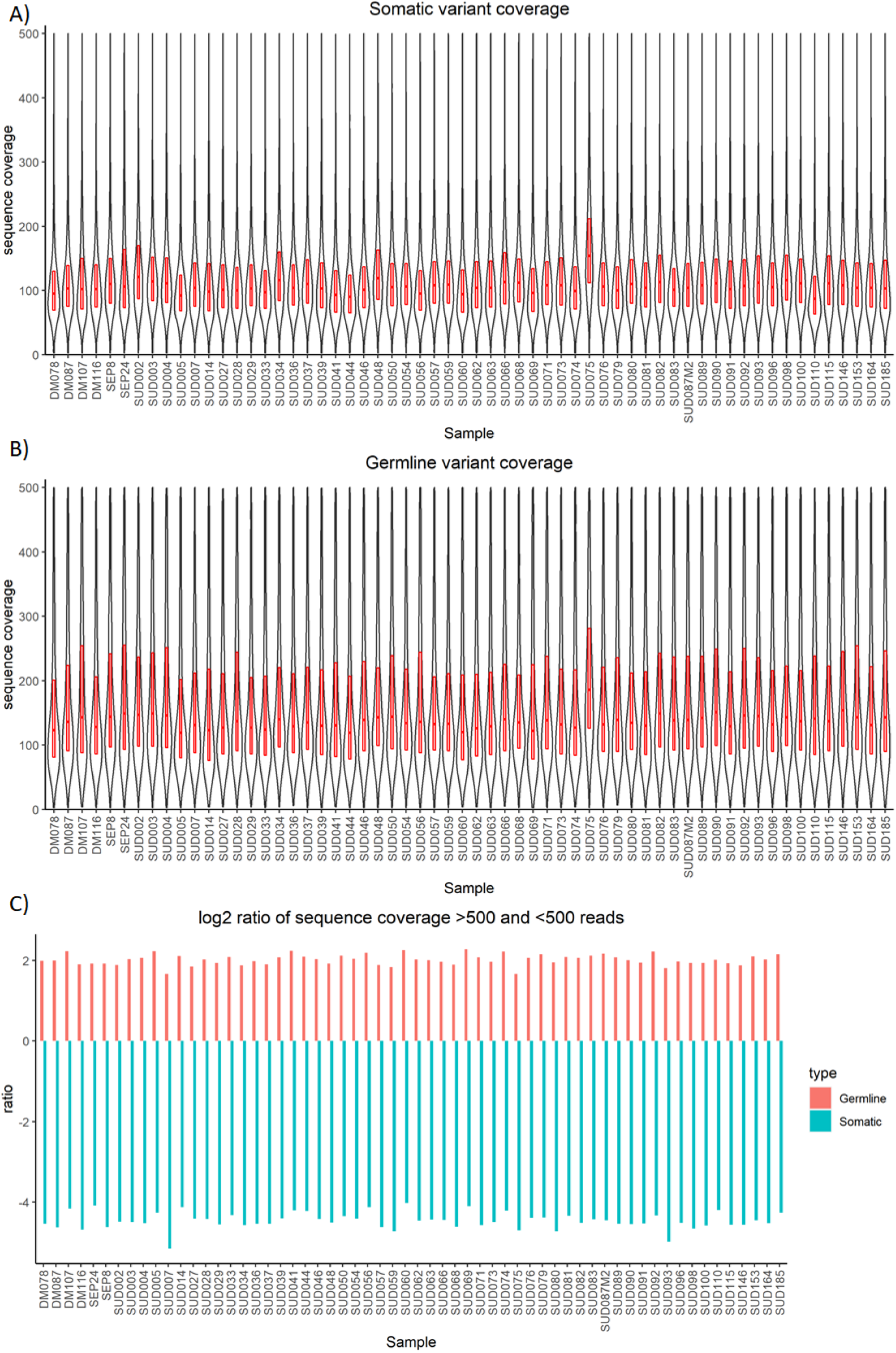
Sequence coverage of somatic (A) and germline (B) mutations below 500 reads. log2 ratio of several variants with more than 500 reads and less than 500 reads is somatic and germline mutations (C) in WES data of 60 tumor samples indicating the lower sequence coverage of somatic mutations compared to the germline.

**Figure S4.**
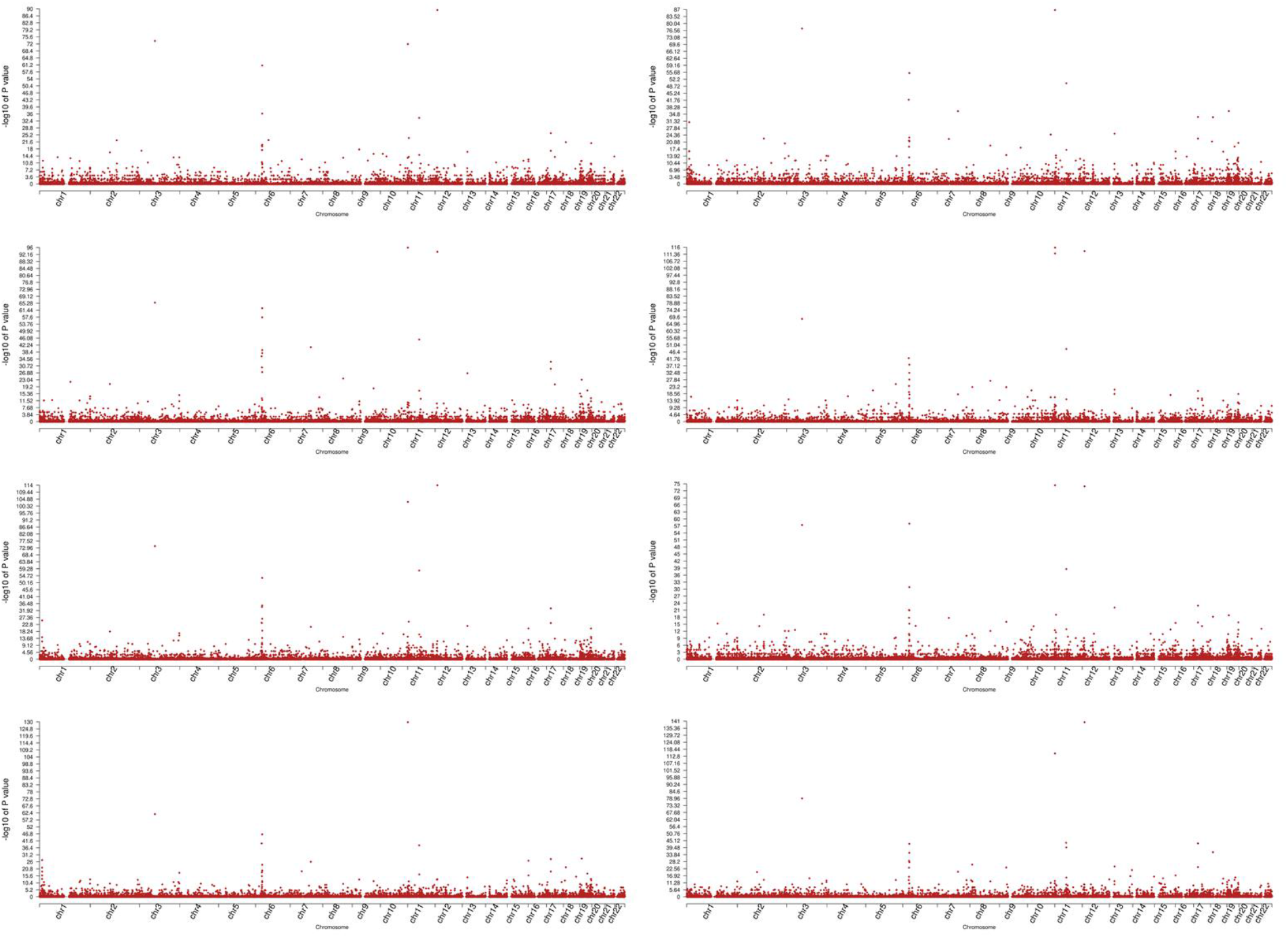
Manhattan plots showing the enrichment of somatic mutations compared to germline in 6 tumor samples shown as examples. Low p-values (I.e., a high −log_10_(p-value)) indicate a high mutation load of genes at a given chromosomal location (x-axis), indicating a hot spot. Genes found to be significant in at least ⅓ of the 60 samples were selected as a hotspot in this study and are listed in Table S3. All sample shows a peak in chromosome 6 corresponding to the HLA region.

**Figure S5.**
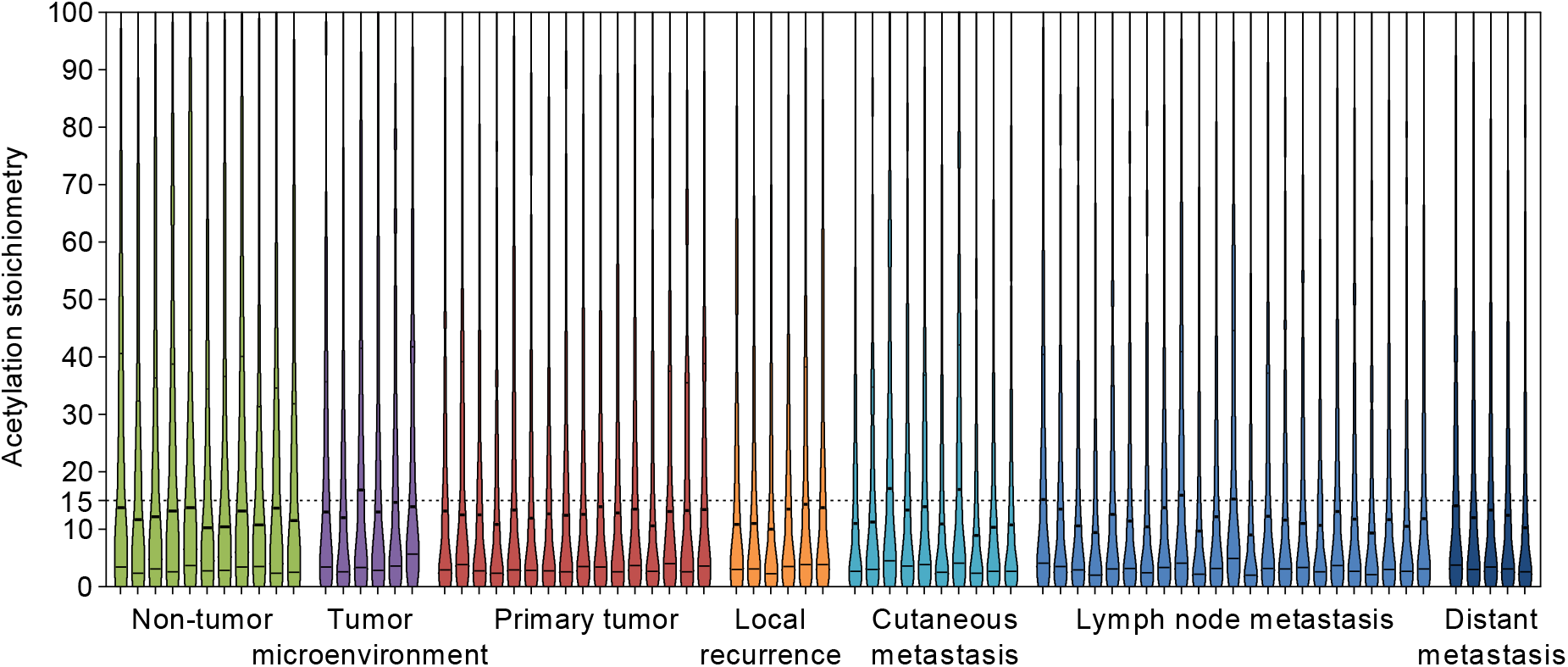
Distribution of the lysine acetylation stoichiometry in the percentage of occupancy of the target site, in all the solid biopsies enrolled in the study. The dashed line indicates a site-specific acetylation occupancy of less than 15%.

## 8. Supplementary Tables

**Table S1.**
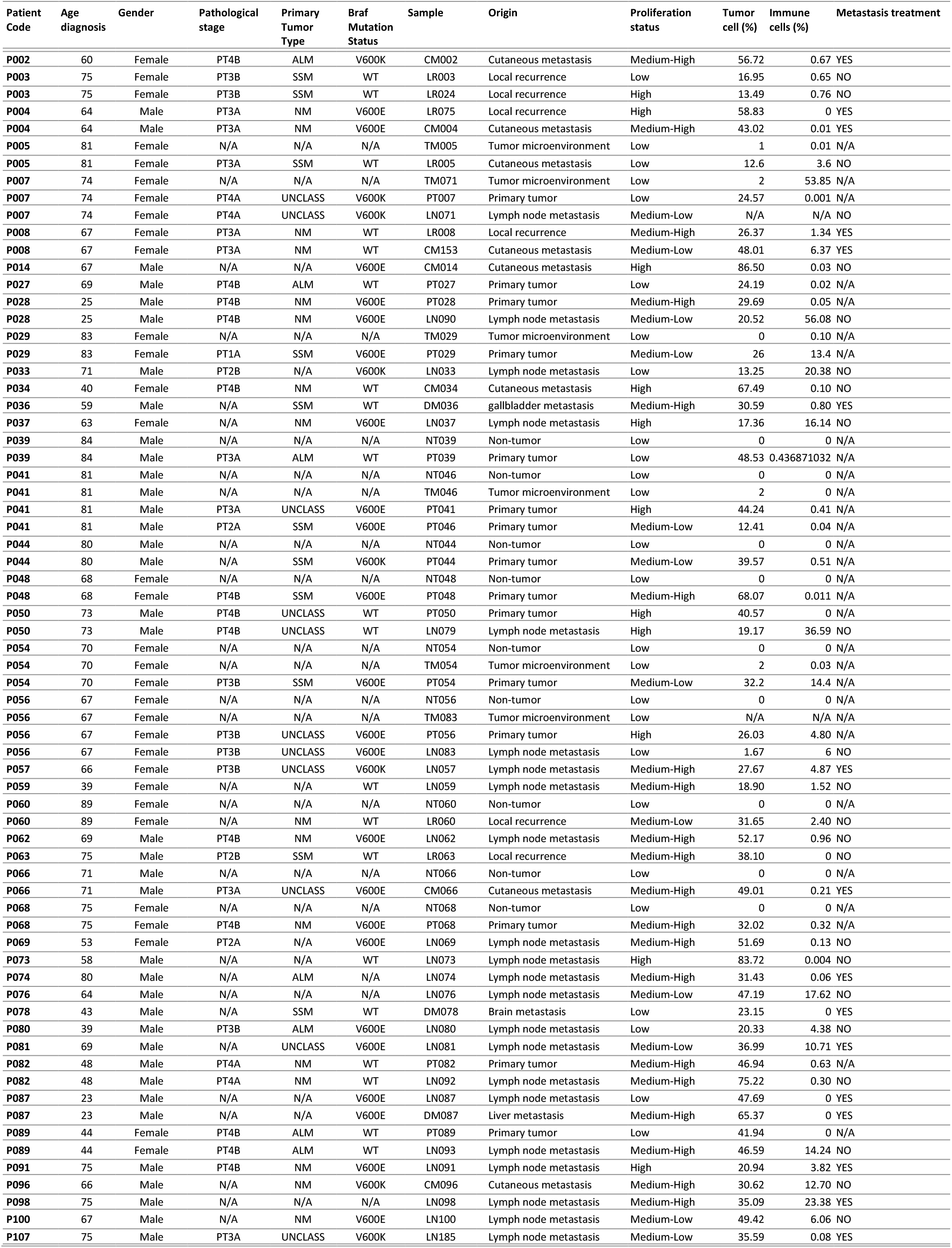
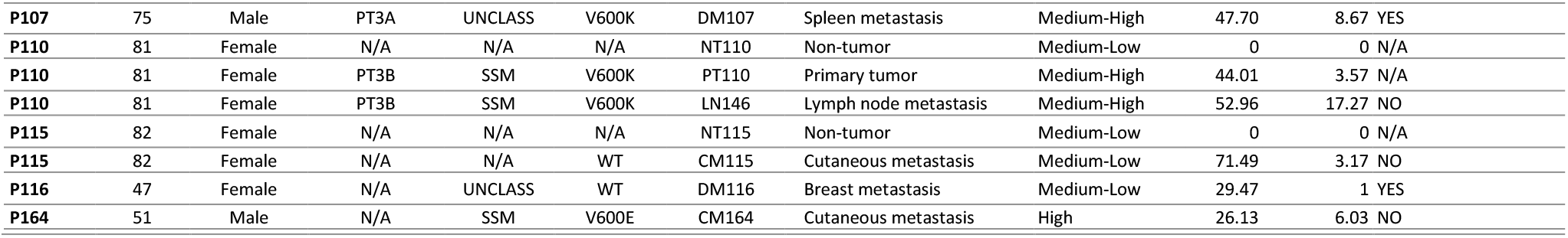
Relevant information of the patients and samples.

**Table S2.** Summary of average (± standard variation) of variants (SNPs, indels) identified in WES data of 60 tumors.

| Type of mutations | Somatic | Germline |
| --- | --- | --- |
| SNPs A>C and T>G | 4361 ± 716 | 872 ± 29 |
| SNPs A>G and T>C | 4573 ± 635 | 6104 ± 119 |
| SNPs A>T and T>A | 11681 ± 1916 | 738 ± 41 |
| SNPs C>A and G>T | 9497 ± 1622 | 1158 ± 62 |
| SNPs C>G and G>C | 5585 ± 888 | 1325 ± 42 |
| SNPs C>T and G>A | 10479 ± 1593 | 8818 ± 299 |
| All SNPs | 46177 ± 7162 | 19015 ± 547 |
| indels | 57423 ± 10369 | 256 ± 19 |
| All variants | 113609 ± 19156 | 19271 ± 256 |

**Table S3.**
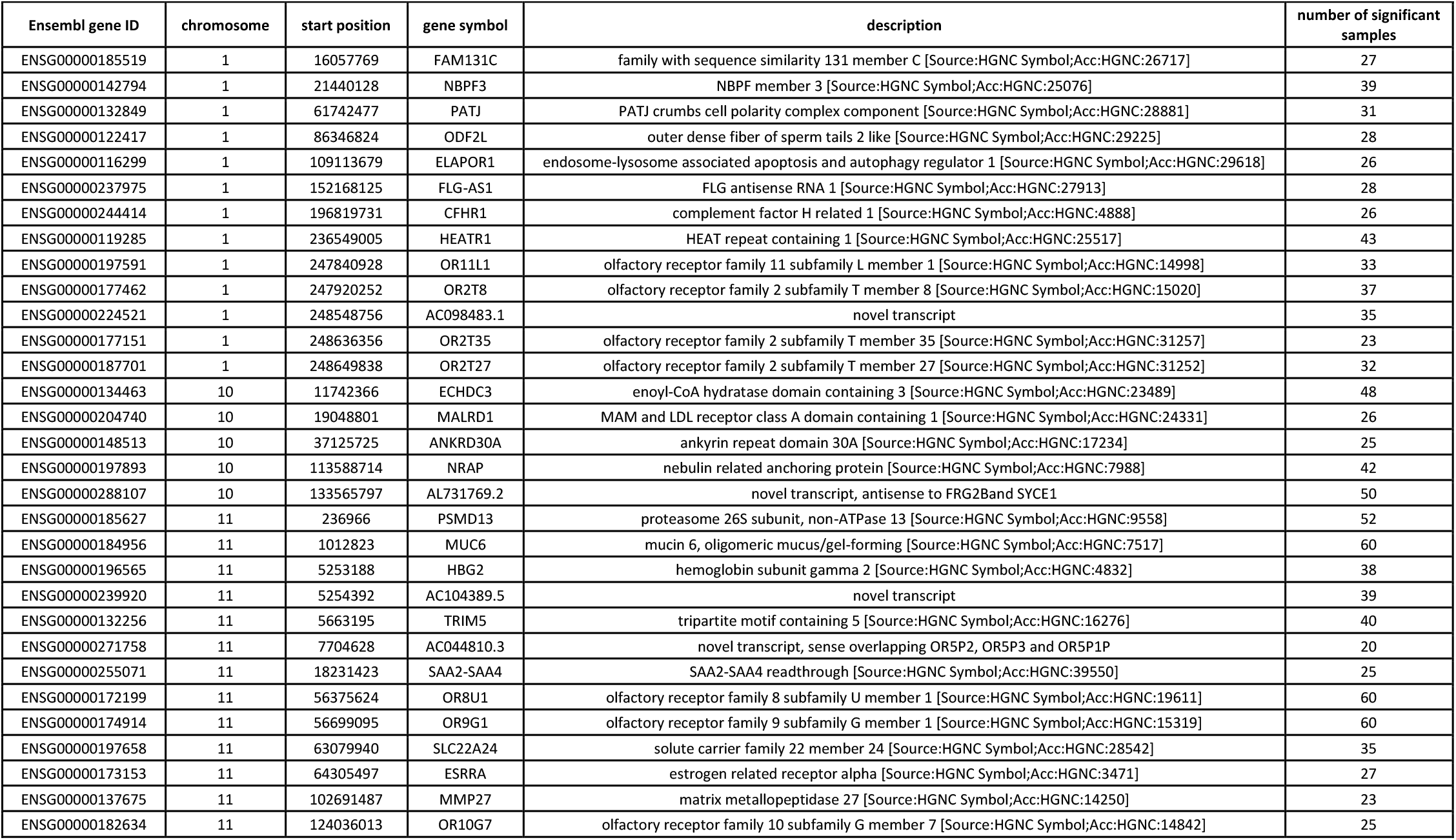
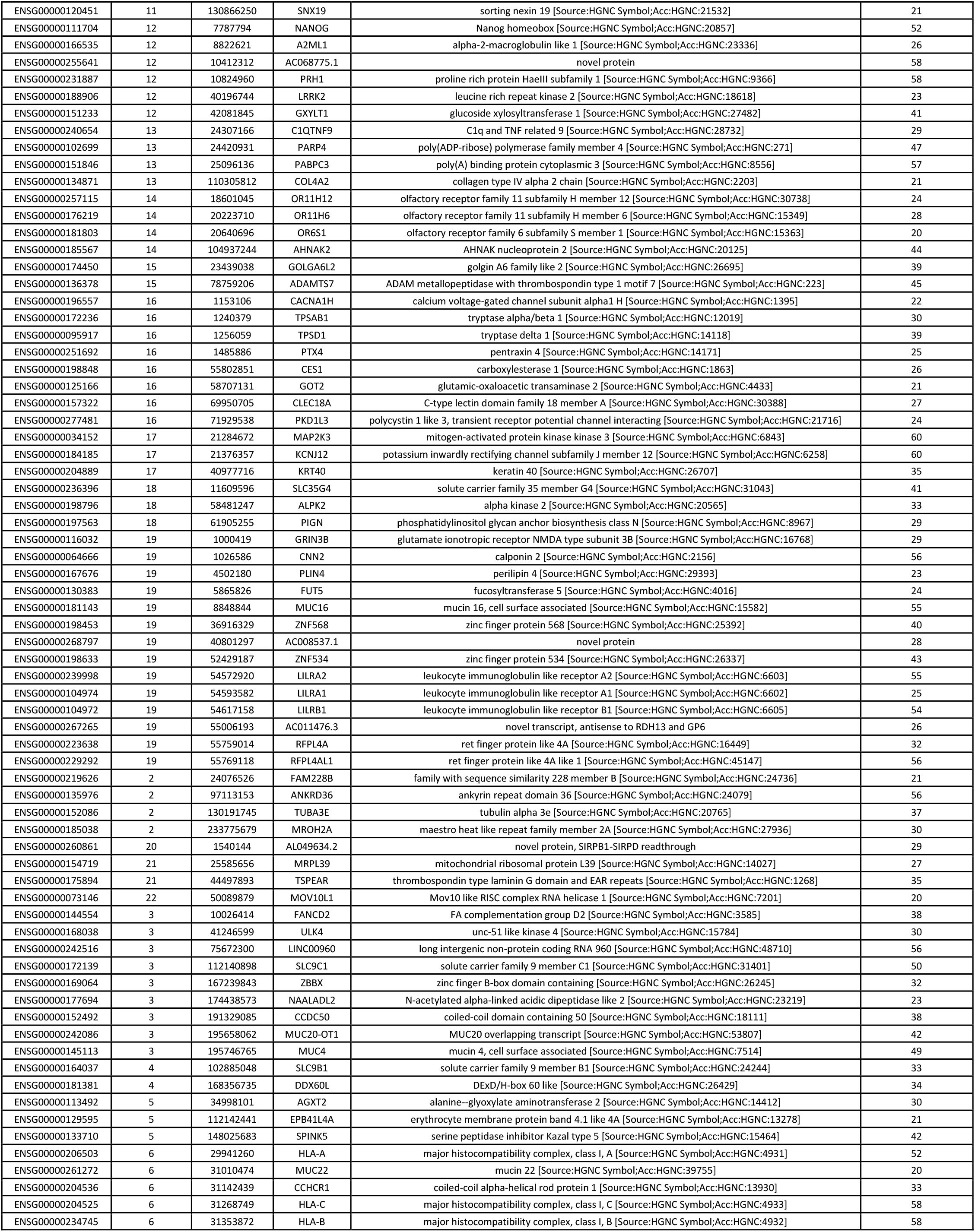
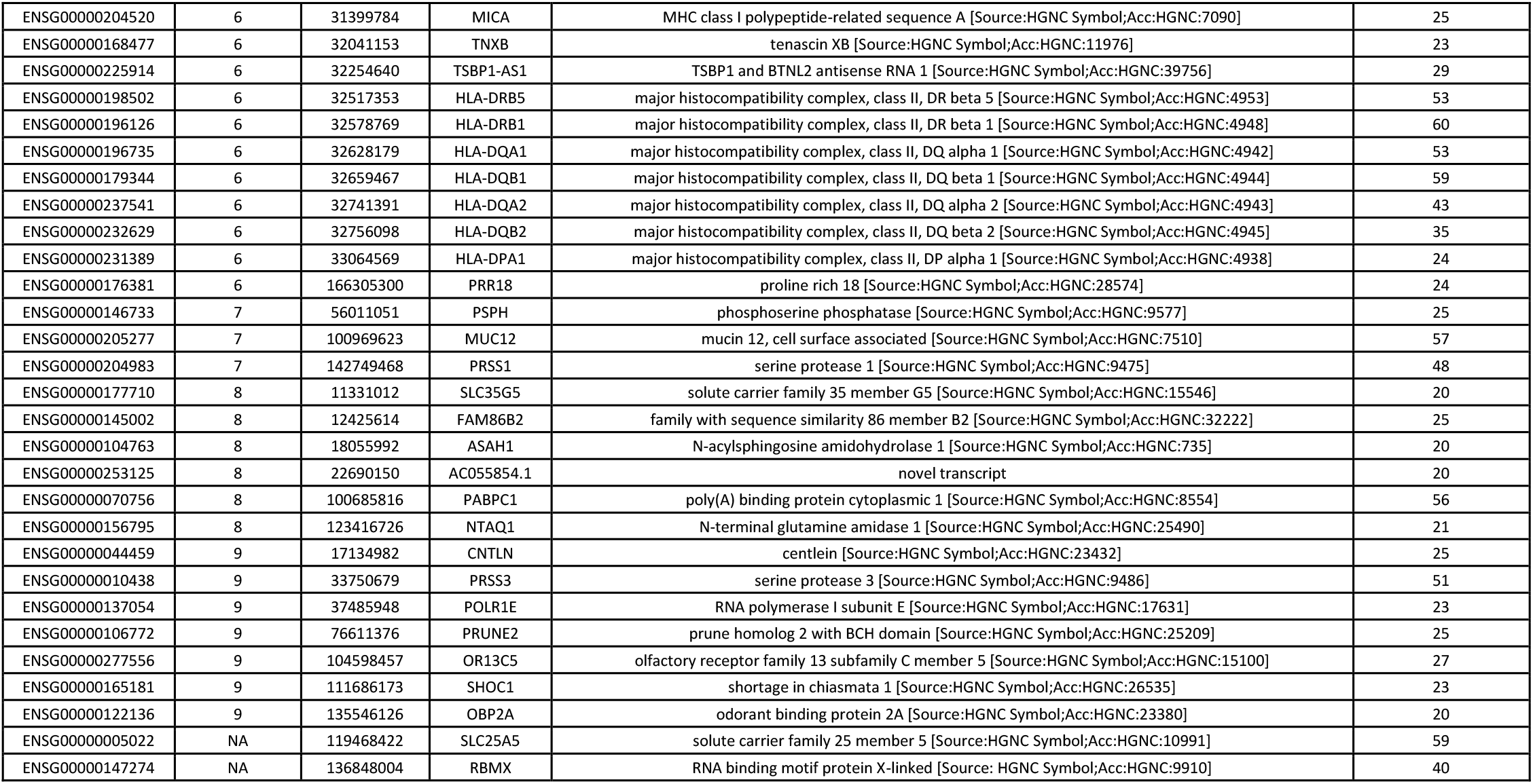
List of genes identified as a hotspot in melanoma WES data requiring presence of the hot spot in 20 out of the 60 analyzed samples.

**Table S4.** Summary statistics showing the average number of somatic and germline mutations in 16 KEGG pathways known to be dysregulated in melanoma. The pathways were selected our proteomic data and previous knowledge.

| No. of genes in the pathway | No. of genes detected in WES data | KEGGID | Pathway | Number of somatic mutations | Number of germline mutations |
| --- | --- | --- | --- | --- | --- |
| 133 | 105 | hsa00190 | Oxidative phosphorylation | 202 ± 52 | 17 ± 4 |
| 51 | 45 | hsa00330 | Arginine and proline metabolism | 187 ± 45 | 21 ± 4 |
| 48 | 46 | hsa00280 | Valine, leucine and isoleucine degradation | 146 ± 41 | 20 ± 3 |
| 57 | 47 | hsa00480 | Glutathione metabolism | 153 ± 38 | 11 ± 3 |
| 151 | 123 | hsa03040 | Spliceosome | 485 ± 117 | 26 ± 7 |
| 98 | 88 | hsa03015 | mRNA surveillance pathway | 422 ± 98 | 56 ± 10 |
| 46 | 42 | hsa03050 | Proteasome | 120 ± 31 | 12 ± 4 |
| 294 | 276 | hsa04010 | MAPK signaling pathway | 1568 ± 313 | 118 ± 13 |
| 240 | 214 | hsa04020 | Calcium signaling pathway | 1578 ± 300 | 179 ± 14 |
| 354 | 317 | hsa04151 | PI3K-Akt signaling pathway | 1932 ± 383 | 222 ± 17 |
| 120 | 109 | hsa04152 | AMPK signaling pathway | 634 ± 131 | 48 ± 7 |
| 166 | 158 | hsa04310 | Wnt signaling pathway | 832 ± 170 | 55 ± 6 |
| 67 | 52 | hsa00010 | Glycolysis / Gluconeogenesis | 218 ± 54 | 28 ± 5 |
| 140 | 131 | hsa04120 | Ubiquitin mediated proteolysis | 701 ± 179 | 45 ± 6 |
| 79 | 75 | hsa03018 | RNA degradation | 315 ± 79 | 55 ± 11 |
| 109 | 95 | hsa04066 | HIF-1 signaling pathway | 494 ± 112 | 35 ± 6 |

